# Bumblebees can detect floral humidity

**DOI:** 10.1101/2021.03.19.436119

**Authors:** Michael J. M. Harrap, Natalie Hempel de Ibarra, Henry D. Knowles, Heather M. Whitney, Sean A. Rands

## Abstract

Floral humidity, a region of elevated humidity proximal to the flower, occurs in many plant species and may add to their multimodal floral displays. So far, the ability to detect and respond to floral humidity cues has been only established for hawkmoths when they locate and extract nectar while hovering in front of some moth-pollinated flowers. To test whether floral humidity can be used by other more widespread generalist pollinators, we designed artificial flowers that presented biologically-relevant levels of humidity similar to those shown by flowering plants. Bumblebees showed a spontaneous preference for flowers which produced higher floral humidity. Furthermore, learning experiments showed that bumblebees are able to use differences in floral humidity to distinguish between rewarding and nonrewarding flowers. Our results indicate that bumblebees are sensitive to different levels of floral humidity. In this way floral humidity can add to the information provided by flowers and could impact pollinator behaviour more significantly than previously thought.

**Summary statement:** We demonstrate for the first time that bumblebees show a preference to elevated floral humidity and can learn to distinguish flowers that differ in floral humidity levels.

## Introduction

Floral humidity, an area of elevated humidity within the headspace of the flower, has been demonstrated to occur in a number of flower species (Corbet et al., 1979; Nordström et al., 2017; von Arx et al., 2012). Floral humidity is created by a combination of nectar evaporation and floral transpiration (Azad et al., 2007; Corbet et al., 1979; Harrap et al., 2020a; von Arx et al., 2012) although the contribution of these two influences may vary between species. Transects of the flower headspace of 42 species found 30 (71%) produce floral humidity of an intensity greater than would be expected from any conflating environmental humidity sources (Harrap et al., 2020a) (such as the minimal humidity differences due to uneven air mixing in the sampling room, or humidity produced by the capped horticultural tubes that flowers were mounted in during sampling). The intensity of floral humidity produced by flowers, represented by 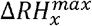 (the average peak difference in relative humidity in the flower species’ headspace, compared to the background), reached up to 3.71% (in *Calystegia sylvatica*). Floral humidity occurs widely and varies between species (Harrap et al., 2020a) and does not appear to be limited to species visited by a particular group of pollinators (Harrap et al., 2020a), elevated floral humidity intensity has been observed in flowers pollinated primarily by moths (von Arx et al., 2012), flies (Nordström et al., 2017) and bees (Corbet et al., 1979).

Whether such variations in floral humidity can be used as a foraging cue is poorly understood, and has only been demonstrated in a single pollinator species, *Hyles lineataIa*, a hawkmoth frequently pollinating *Oenothera caespitosa* (von Arx, 2013; von Arx et al., 2012). It was demonstrated that *H. lineatala* shows a preference to artificial flowers producing floral humidity comparable to that produced by *O. caespitosa*, over those at ambient humidity. Investigation of the capacity of pollinators other than *H. lineatala* to respond to floral humidity is limited (von Arx, 2013), with non-experimental observations that flies may use floral humidity in addition to other floral display traits produced within Indian alpine environments (Nordström et al., 2017). Given that floral humidity is present in many flower species, as recently measured by Harrap et al. (Harrap et al., 2020a), it is most likely that floral humidity is regularly encountered as part of flowers’ multimodal displays by a wide range of generalist pollinators and influences their foraging behaviours.

Sensitivity to environmental (non-floral) humidity is well reported in insects (Enjin, 2017; Havukkala and Kennedy, 1984; Kwon and Saeed, 2003; McCall and Primack, 1992; Peat and Goulson, 2005). Honeybees *Apis mellifera* respond to humidity levels within the nest, regulating humidity to different levels in different parts of the nest (Human et al., 2006; Nicolson, 2009). Elevated humidity triggers nest ventilation behaviours in bees such as fanning nest structures, and low humidity encourages behaviours that increase nest humidity by the evaporation of nectar water or water collection (Abou-Shaara et al., 2017; Ellis et al., 2008; Human et al., 2006). Biting flies and mosquitoes are thought to respond to humidity given off by their host organisms, among other cues (Chappuis et al., 2013; Olanga et al., 2010; Smart and Brown, 1956). Mosquitoes also make use of humidity to locate still-water oviposition sites (Okal et al., 2013). Furthermore, following the presentation of sugar water droplets that touch their antenna, restrained honeybees have been seen to show a proboscis extension response to droplets of water placed near (but not touching) the antenna (Blenau and Erber, 1998; Kuwabara, 1957; Mercer and Menzel, 1982). Such responses are likely in response to the water vapour (= humidity) given off by the droplet, suggesting that bees can be conditioned based on humidity to some degree (Blenau and Erber, 1998; Kuwabara, 1957; Mercer and Menzel, 1982). Taken together with the presence of hygrosensitive (humidity detecting) sensilla in many pollinating insects, this suggests that pollinator groups other than hawkmoths possess the necessary sensory mechanisms to detect and respond to humidity cues and signals in the context of foraging on flowers.

The presence of a hygrosensitive antennal sensillum, the ceolocapitular sensillum (Yokohari, 1983; Yokohari et al., 1982), has been reported for bees (Ahmed et al., 2015; Tichy and Kallina, 2014; Yokohari, 1983; Yokohari et al., 1982); these sensilla are common and show a wide distribution across the antenna in *Bombus* bumblebees (Fialho et al., 2014). This may allow bumblebees to show higher humidity sensitivity (Fialho et al., 2014), although the exact mechanism by which these ceolocapitular sensilla detect humidity is uncertain (Enjin, 2017; Tichy and Kallina, 2010). Insects always possess two types of humidity-sensitive cells within ceolocapitular sensilla: dry cells, which respond to a lack of humidity; and moist cells, that respond to its presence (Yokohari, 1983; Yokohari et al., 1982). In addition to signalling based on the humidity at a given instant, moist and dry cells signal with at a greater frequency in response to the rate of humidity changes (Tichy, 2003; Tichy and Kallina, 2014, 2010). Insects can therefore detect both the humidity at a given time, and also the rate and direction of humidity changes, getting drier or moister. The sensitivity to humidity reported in *Apis mellifera* (Tichy and Kallina, 2014) echoes the humidity changes produced by flowers (Harrap et al., 2020a), suggesting that floral humidity differences could feasibly be detected by these pollinators, allowing them to use floral humidity while foraging.

We investigated the capacity of bumblebees *Bombus terrestris* to detect and respond to artificial flowers producing floral humidity at levels comparable to the floral humidity detected in real flowers. Furthermore, in these experiments we explored the nature of bumblebees’ responses to floral humidity, asking whether floral humidity cues influence their spontaneous flower choices. As pollinators can associate differences between flowers in various floral traits to distinguish more rewarding flowers from less rewarding flowers, such as colour (Streinzer et al., 2009), scent (Daly et al., 2001; Galen and Newport, 1988), floral temperature (Dyer et al., 2006; Whitney et al., 2008), floral texture (Kevan and Lane, 1985), electrostatic properties (Clarke et al., 2013) and patterning of these signals (Harrap et al., 2020b, 2017; Hempel de Ibarra et al., 2015; Lawson et al., 2018; Whitney et al., 2009), we also conducted differential conditioning experiments to investigate the capacity of bumblebees to learn differences in floral humidity and use floral humidity for flower species recognition.

## Materials and Methods

Responses to floral humidity were tested in lab conditions using captive (female worker) bumblebees, *Bombus terrestris audax* (Harris 1776) obtained from Biobest (Westerlo, Belgium via Agralan Swindon, UK). Bumblebees are an appropriate choice of forager to test responses to floral humidity as they visit a wide range of species, including many of those found to produce different levels of floral humidity by Harrap et al. (Harrap et al., 2020a). For example, bee pollinators are known to forage on species throughout the range of floral humidity observed (Harrap et al., 2020a), such as 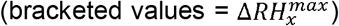: *Calystegia sylvatica* (3.71%); *Eschscholzia californica* (3.24%); *Scabiosa* (1.36%); *Osteospermum* (1.20%); *Papaver cambricum* (0.29%); *P. rhoeas* (0.29%); and *Fuchsia* (0.05). Bee husbandry, marking for identification and the flight arena are described in detail elsewhere (Harrap et al., 2020b, 2019, 2017; Lawson et al., 2018; Pearce et al., 2017). No ethical permissions were required for the experiments involving bumblebees, but the experiments were conducted according to ASAB/ABS guidelines.

We created two types of artificial flower that produced artificial floral humidity comparable in intensity and structure observed on natural flowers. In ‘active’ flowers, elevated humidity was created by pumping humid air into the flower, whereas in ‘passive’ flowers, a wettable sponge was placed within the flower to generate elevated floral humidity. The two types of artificial flowers allowed us to observe whether responses of bees were the same independently of how floral humidity was generated. The active flowers echoed the hawkmoth-flower design used by von Arx et al. (von Arx et al., 2012), but were adapted to better suit bumblebee foraging behaviour. In this way active flowers allowed us to test bee responses to a stimulus produced in a manner comparable to that study.

Both types of flowers had two variants that varied in the level of humidity they produced. The ‘humid’ variant produced elevated humidity in the proximity of the flower’s top, whilst in a ‘dry’ artificial flower variant floral humidity was much lower. Full details of both artificial flower designs and maintenance are given in Supporting Information Appendix 1.

To reward the bees, a well (created from the lid of an Eppendorf tube) in the centre of the artificial flowers provided a drop of sugar solution in rewarding flowers, and a drop of water in non-rewarding flowers. All artificial flowers were dry to the touch (to avoid conflating responses to wet flower surfaces) and did not differ in temperature or other characteristics bees could respond to.

Floral humidity produced by artificial flowers, both variants (humid and dry) of both type (active and passive), was measured and analysed using the procedures described in Supporting Information Appendix 2. As liquid in feeding wells of flowers (water or sucrose solution) might produce humidity, artificial flowers were sampled with a 25*µ*l droplet of water in their feeding wells, which meant that we were able to assess the humidity produced by artificial flowers as they are presented to the bees (see below). This measurement and analysis of floral humidity mirror those used by Harrap et al. (Harrap et al., 2020a), allowing direct comparison of floral humidity between artificial and natural flowers.Floral humidity levels produced by both artificial flower types was comparable to that produced by ‘real’ natural flowers (Harrap et al., 2020a). The humidity intensity (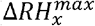, the average peak difference in relative humidity in the flower species’ headspace, compared to the background) of both active and passive humid flower variants was 3.08% and 3.49% respectively, and was comparable to flower species that produce higher intensity floral humidity, such as: *Calystegia sylvatica* (3.71%); *Eschscholzia californica* (3.24%), *Taraxacum* agg. (3.35%), or *Ranunculus acris* (3.41%). Humidity intensity produced by dry artificial flower variants was more comparable to intermediate levels of humidity observed on flowers. Active dry flowers had a 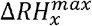 of 0.92%, similar to intensities seen in *Convolvulus sabatius* (0.87%), *Cyanus segetum* (1.10%) and *Linum usitatissium* (0.8%). Passive dry flowers had a 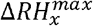 of 2.13%, while not matching the intensity of specific species this is in a similar range to that of *Leucanthemum vulgare* (1.79%) and *Achillea millefolium* (1.73%), and well within the range of floral humidity intensity observed.

Floral humidity structure was similar to that of natural flowers ((Harrap et al., 2020a) and Figures S3 and S4 in supplementary file 1, appendix 2), showing a peak value in the centre of the flower and declining with distance from the flower. Lastly, when refilling of feeding wells and passive flower internal components in bee trials (see below) was considered, artificial floral humidity appeared to remain largely stable across the timescales of bee trial experiments. Consequently, differences in humidity between humid and dry variants remained at ∼2.16% for active flowers and ∼1.36% for passive flowers. Therefore, the artificial floral humidity presented to bumblebees in the bee trials remained comparable to natural flowers, in terms of floral humidity intensity, structure and differences between flowers. For further detail on artificial floral humidity produced by artificial flowers see supplementary file 1, appendix 2.

### Bee trials

Two kinds of experiments were carried out on captive bumblebees. Firstly, preference experiments (*e*.*g*. see (Dyer et al., 2006; Lehrer et al., 1995; von Arx et al., 2012)) were carried out using both artificial flower types. Secondly, differential conditioning techniques (*e*.*g*. see (Clarke et al., 2013; Dyer and Chittka, 2004; Harrap et al., 2017; Lawson et al., 2018)) were carried out with passive artificial flowers only. This was due to the limits on how much and how quickly active artificial flowers could be moved about the arena due to their piping (see construction and experimental details in Supporting Information Appendix 1). Both preference and learning (differential conditioning) experiments test the capacity of bees to detect and respond to floral humidity differences. Additionally, preference experiments investigate how differences in floral humidity between flowers, in the absence of any other differences, may influence flower choice of naïve bees with no previous experience of floral humidity. Learning experiments (differential conditioning) investigate whether bees can associate differences in floral humidity with corresponding differences in rewards and use this to inform foraging choices. Therefore, two experiments together assess different ways that floral humidity might influence the foraging behaviours bees. Individual bees were not reused between experiments: an individual bee would only take part in one experiment (preference or conditioning) as part of a single test group (see below) or on a single type of artificial flower (active or passive).

### Preference experiments

In preference experiments, individual bees had their preference for floral humidity tested using either active or passive flowers. Two different bumblebee nests were used in the passive flower tests. Bees used in the active flower tests came from four different nests, which included the two nests used in the passive flower tests. During preference tests, bees were presented with eight artificial flowers of the type assigned to them, placed randomly about the foraging arena floor. Four of these were the humid flower variant, and the other four were the dry flower variant. All artificial flowers were rewarding, containing a 25*µ*l droplet of 30% sucrose solution within their feeding wells.

Individual bees were released into the arena alone, and bees were allowed to forage freely on these artificial flowers, and were free to return to the nest at all times. We monitored whether bees made contact with the top of artificial flowers (which was recorded as a landing behaviour), and whether the bee extended its proboscis into the feeding well (‘fed’) or left without doing so at each landing. After a bee had fed on a flower, the flower was refilled and moved. Bees are highly capable of learning the locations of rewarding flowers (Burns and Thomson, 2006; Robert et al., 2017), particularly within the small area of a flight arena. It would be possible for bees to learn to return to the locations where rewards had been found previously. Therefore, as conducted by previous studies (Harrap et al., 2019, 2017), we changed flower position after a bee fed to reduce the possibility of bees using spatial learning to inform foraging decisions. In the passive flowers this involved taking the flower out of the arena and placing it back in in a different position. With active flowers, the ability to move the flowers was limited by their pipes and the flowers’ current arena entry points. Consequently, active artificial flowers were not taken out of the arena but were instead moved to a different point. In the rare instances where a bee fed from a flower and revisited it before it could be moved, then these revisits were not counted. When the bee returned to the nest all the flowers were removed from the arena, cleaned and returned to the arena in a new position as described in Supporting Information Appendix 1.

For each test bee, this cycle of moving flowers continued until the bee had made at least 20 flower landings. This was normally achieved in 4.38 ± 0.44 (mean ± SEM) foraging bouts (a ‘foraging bout’ being the time between departure and return to the nest) with an average of 5.06 ± 0.41 visits per bout for bees presented with active flowers and 3.00 ± 0.34 bouts with an average of 7.40 ± 0.67 visits per bout for those presented with passive flowers. Sixteen bees completed the preference experiment on each type of artificial flower (32 bees in total).

The rate at which bees made a positive response to floral humidity (the ‘humidity response rate’) over the 20 flower landings was calculated for each bee. A positive response was classed as either landing on a humid flower and extending the proboscis into the feeding well, or landing on a dry flower and leaving without extending the proboscis into the feeding well. The humidity response rate data was bounded between 0 and 1, and so was arcsine square root transformed to fit test assumptions. We used a two-tailed Wilcoxon signed-rank test to test whether the median value of the transformed humidity response rate differed from that expected from random choice (a 0.5 humidity response rate, 0.79 once arcsine square-root transformed), using *R* 3.6.3 (R Development Core Team, 2017).

Temperature differences between humid or dry flower variants might occur as a result of evaporative water loss or action of mechanical components within artificial flowers. As bees can respond to floral temperature showing preferences and learning (Dyer et al., 2006; Harrap et al., 2020b, 2017; Whitney et al., 2008), differences above the temperature sensitivity of bees may conflate bee responses. For this reason, flower temperature differences of artificial flowers were monitored alongside the preference experiment, using a thermal camera (FLIR E60bx, FLIR systems, Wilsonville, USA), to see if the flowers develop a temperature difference bees could respond to (a temperature difference greater than 2°C (Heran, 1952)). This was done at the start of foraging or after flower cleaning at the end of foraging bouts, by randomly selecting one humid and one dry artificial flower and measuring the temperature of the flower top. Emissivity parameter value used was 0.95, an accepted value for plastics (Harrap et al., 2018), and reflected temperature used was a consistent value of 20°C.

### Learning experiments

In learning experiments only the passive artificial flowers were used. Individual bees were allowed into the arena alone and were presented with eight artificial flowers placed randomly about the flight arena: four rewarding and four nonrewarding with humidity production by these flowers assigned as per the bee’s test group. Bees were allowed to forage freely on these artificial flowers, again allowed to return to the nest as required. Whether bees made contact with the top of artificial flowers, classed as landing, was monitored. Whether a bee extended its proboscis into the feeding well or left without doing so at each landing was also monitored.

Before a trial began, bees were assigned to one of three test groups: ‘Humid rewards’ group, where the humid passive flower were rewarding and the dry passive flowers non-rewarding; *ii*) ‘Dry rewards’ group, where the rewarding flowers were dry passive flowers, and humid passive flowers were non-rewarding; *iii*) ‘Control’ group, where none of the flowers produced humidity (*i*.*e*. both rewarding and nonrewarding were dry passive flowers), meaning that flowers only differed in their rewards. Rewarding flowers had a 25*µ*l droplet of 30% sucrose solution within their feeding wells and nonrewarding flowers contained a 25*µ*l droplet of water. Four different bumblebee nests were used in this experiment, none of these nests were used in preference experiments.

The ‘Control’ group was required for checking to what extent bees could use any miscellaneous cues other than humidity or variables present in the experimental setup to solve the task. Such miscellaneous cues may include the shapes of the numbers on the side of the flowers (although effort was made to reduce this possibility, see supplementary file 1, appendix 1), or small differences in the shape of the cut away components of the flowers’ lids (see supplementary file 1, appendix 1) that may influence the appearance or tactile properties of individual flowers. The arrangement of flowers in the arena, such as incidental clustering of rewarding or nonrewarding flowers, or some combinations of environmental cues within the arena may also facilitate learning in some manner. These miscellaneous variables might give rise to basic capacity of bees to find individual rewarding flowers, independently of humidity differences, within the specific settings of our setup.

Bees were observed for 70 flower visits which is well beyond the number of visits needed for bees to learn a salient cue, and sufficient to demonstrate such learning by a consistent change in foraging choices (*e*.*g*. see (Clarke et al., 2013; Harrap et al., 2017; Lawson et al., 2018)). Bees achieved 70 visits in, on average, 5.13 ± 0.31 foraging bouts (mean ± SEM) with 13.78 ± 0.56 landings in each bout. 15 bees completed this learning trial in each test group (45 bees in total).

After a bee fed on a flower and flew off into another part of the arena, that flower was carefully removed from the arena through side openings and refilled with sucrose or water, as appropriate, before being placed back at a different location. This reduced the chance of bees’ associating particular spatial locations with the reward. If a bee suddenly revisited the flower before it could be moved, then these revisits were not counted. When the bee returned to the nest, all the flowers were removed from the arena, cleaned and returned to the arena as described in Supporting Information Appendix 1.

Flower visits were determined as ‘correct’ or ‘incorrect’ using the same criteria described in our previous studies Harrap et al. (Harrap et al., 2019, 2017). A bee was recorded as making a correct decision if she landed on a rewarding flower and extended her proboscis into the feeding well, or if she did not extend her proboscis into the feeding well after landing on a non-rewarding flower. Correspondingly, a bee was recorded as making an incorrect decision if she landed on a non-rewarding flower and extended her proboscis into the feeding well, or did not extend her proboscis into the feeding well when she landed on a rewarding flower. The success rate, defined as the proportion of correct visits over the previous ten visits, was calculated at ten visit intervals (10 visits, 20, 30… *etc*.) for each bee. As it was bounded between 0 and 1, the success rate data were arcsine square root transformed to fit test assumptions. Generalised linear models were fitted to this data using *R*, and AIC model simplification techniques (Richards, 2008) were used to analyse the effects experience on the flowers (number of visits made) and test group (the presence of floral humidity differences between rewarding and nonrewarding flowers) had on bumblebee foraging success. The full detail of the models used for analysis and AIC model simplification procedure is given in supporting information Appendix 3.

## Results

### Artificial flower temperature differences

Artificial flower temperature differences, as measured during the preference experiments, were negligibly small. Dry passive flowers had a temperature that was 0.31 ± 0.03°C (mean ± SEM) higher than in humid passive flowers throughout the experiment. In active artificial flowers, flowers of the humid and dry variants differed even less in temperature. Dry active flowers were 0.03 ± 0.03°C colder than humid active flowers. Measured differences between dry and humid flower variants (dry flower variant temperature minus humid flower variant temperature) ranged from -0.2 to 0.9°C in passive flowers and -0.5 to 0.5°C in active flowers. These differences in temperature were below the threshold of temperature detection by bumblebees (Heran, 1952) and are unlikely to elicit a response by bumblebees.

### Bee trials

In preference experiments, bumblebees showed a higher spontaneous preference for humid flowers when they were allowed to freely choose between four humid and four dry flowers providing sucrose solution (figure 1). The median humidity response rates differed significantly from that expected from random foraging (0.5), both in tests with passive flowers (Wilcoxon Test, *W* = 109 *n* = 16, *p* = 0.006) and active flowers (Wilcoxon Test, *W* = 119, *n* = 16, *p* = 0.001). The median bee humidity response rates in both preference tests were greater than 0.5 (figure 1).

**Figure 1.**
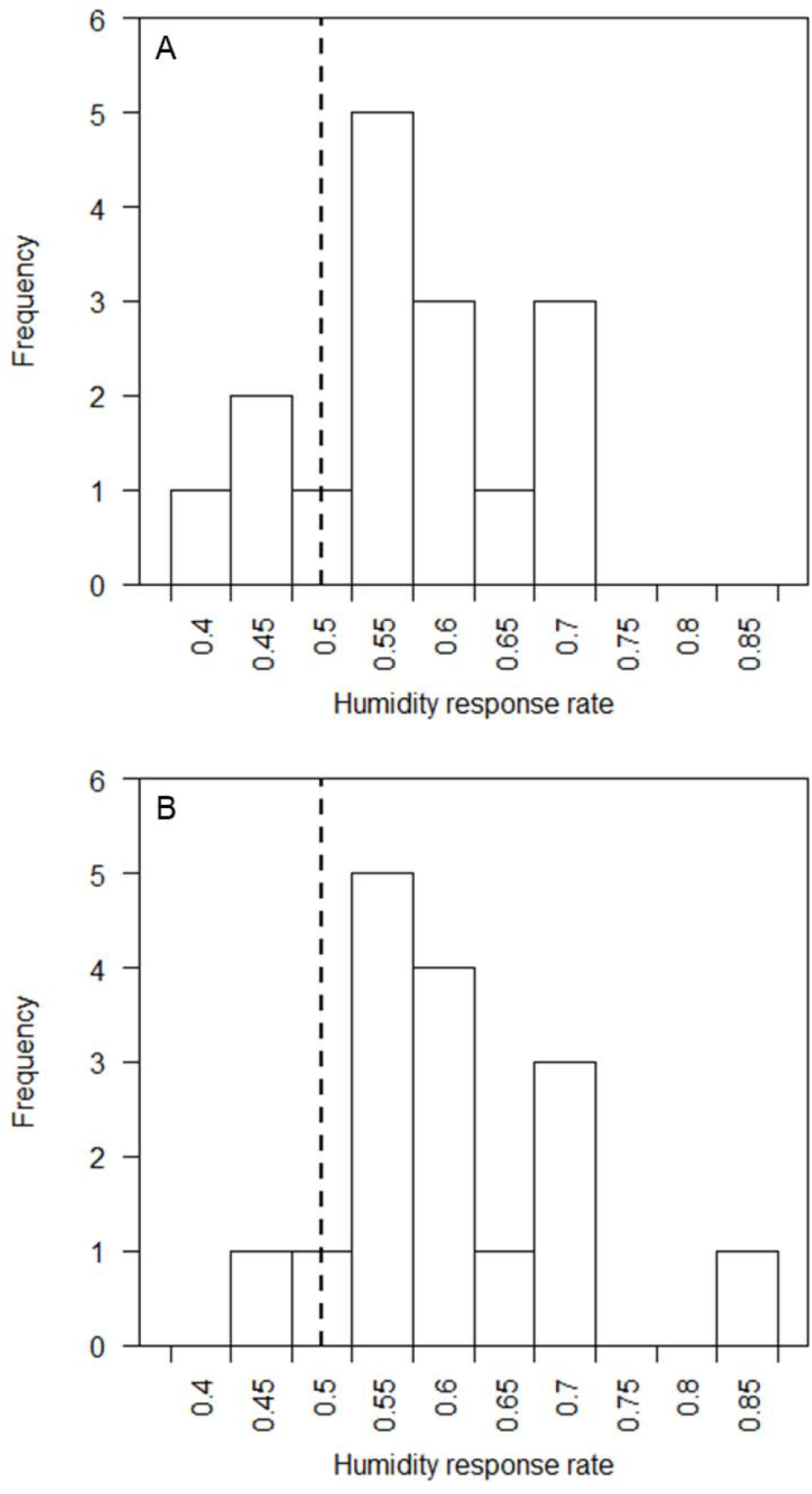
Histograms showing the responses of bumblebees to rewarding passive (**A**) and active (**B**) flowers in the preference experiments. Bars represent the number of bees (*n* = 16 bees in each trial), that over 20 flower landings, achieved each humidity response rate (the proportion of positive responses to ‘humid’ flowers with elevated humidity as opposed to positive responses to ‘dry’ flowers with lower humidity). Dashed vertical line indicates the expected humidity response rate for randomly foraging bees (0.5).

In the learning experiment, bumblebees were presented with passive flowers differing in rewards, providing either sucrose solution or water in feeding wells. The humidity cues corresponding with rewards varied dependent on the three test groups bees were assigned to (see above). The relationship between foraging success (probing the feeding wells of rewarding flowers, or not probing it on nonrewarding flowers) and the experience bees had of the flowers (number of flower visits the bees made) was compared between the three test groups to evaluate the capacity of bumblebees to learn the identify of rewarding flowers based on humidity differences.

The presence of floral humidity differences between rewarding and nonrewarding flowers influenced the ability of bumblebees to learn the identify of rewarding flowers and the foraging success achieved by bees (figure 2). Models that allowed individual bees to have different intercepts as well as different learning speeds independent of their test group (models which had random slopes and intercepts), were not a better fit than those that only allowed individual variation in intercepts (AIC: random slopes and intercepts -298.3 *vs*. random intercepts only -301.3, ΔAIC = 3, Δdeviance = 1.00, df = 2, *p* = 0.605). Models that allowed the number of flower visits made by the bee (experience) to have different influences on success rate between test groups, interacting effects, had a lower AIC (AIC: interaction model -301.29 *vs*. no interaction model -288.16, ΔAIC = 13.13) and a significantly better fit (Δdeviance = 17.13, df = 2, *p* < 0.001) than models that forced experience to have the same effect across test groups. Bees in the control group, where artificial flowers produced no humidity differences, began the experiment with a success rate of 0.5, and improved only slightly, maintaining a success rate just above random over the rest of the experiment. Consequently, models that allowed success of control group bees to change with experience were not better than those that allowed no change in success for control group bees (AIC: experience effects -124.45 *vs*. no experience effects -121.65, ΔAIC = 2.8) although these models were a better fit (Δdeviance = 4.79, df = 1, *p* = 0.029).

**Figure 2.**
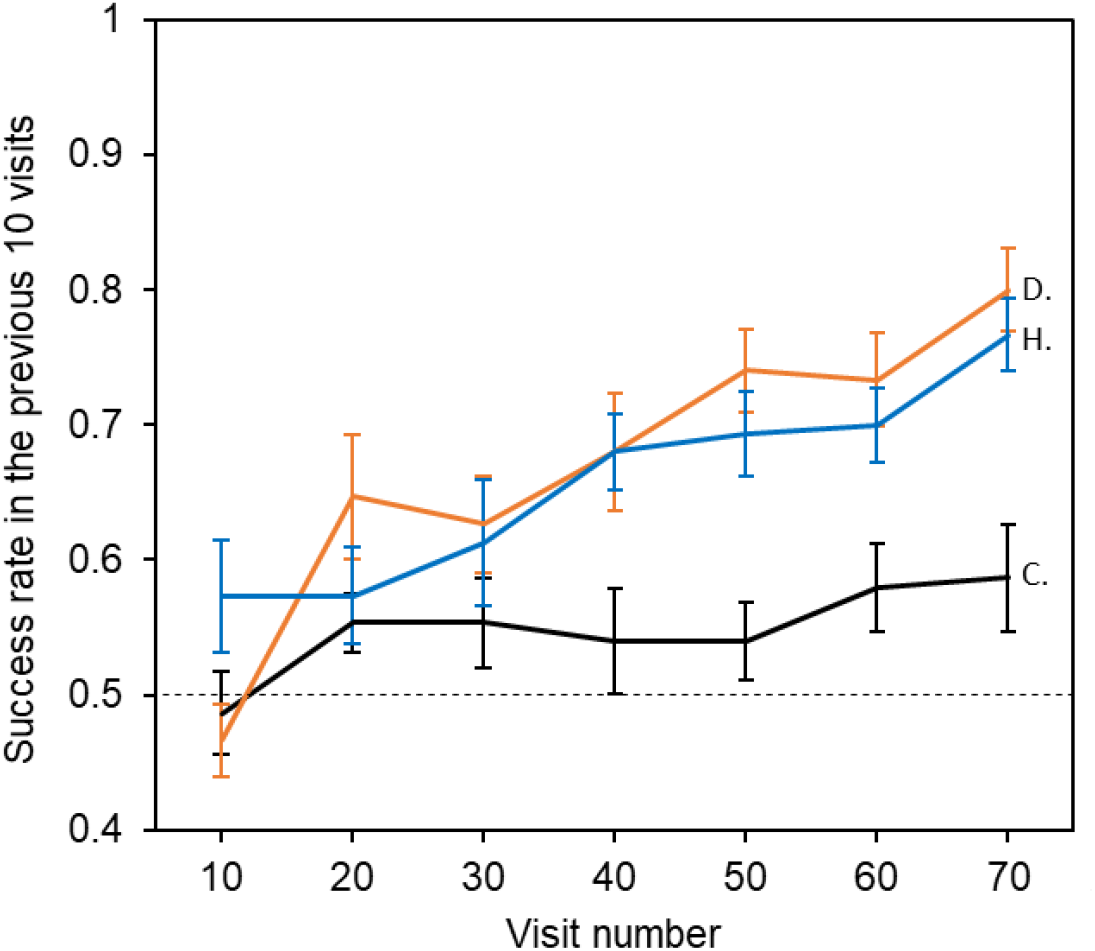
The relationship between bees’ foraging success and experience of passive artificial flowers (flower visits the bee made). Dotted line indicates the 50% success level. Solid lines indicate the mean foraging success of bees in the previous 10 visits. Error bars represent ±SEM. Colour and label of solid lines and error bars correspond with test group: black, the control group, labelled ‘C’.; orange, Dry rewards group, labelled ‘D.’; blue, Humid reward group labelled ‘H’. Number of bees in each test group *n* = 15.

In test groups where floral humidity differed between rewarding and nonrewarding flowers, bees began favouring humidity-producing flowers as naïve bees had in the preference test. In both these test groups, success improved as bees made more flower visits. Bees in both the humid reward and dry reward test groups reached similar success rates by the end of the experiment and both achieved a greater level of success than bees in the control group (figure 2). Models that allowed success rate to change with experience had lower AIC and better fit for both the humid rewards test group (AIC: experience effects -100.0 *vs*. no experience effects -79.7, ΔAIC = 20.24; Δdeviance = 22.24, df = 1, *p* < 0.001) and the dry rewards test group (AIC: experience effects -74.8 *vs*. no experience effects - 36.7, ΔAIC = 38.16; Δdeviance = 40.16, df = 1, *p* < 0.001). This indicates that floral humidity differences improved bumblebee foraging success and learning of the identity of rewarding flowers, regardless of whether rewarding flowers produced higher or lower floral humidity intensities.

## Discussion

By experimentally varying the levels of floral humidity in artificial flowers within a range that is biologically relevant, we show here that bumblebees are able to detect and utilise humidity differences in a flower foraging context. In an all-rewarding array with artificial flowers that offered low and high floral humidity cues, bumblebees showed an unlearned preference for flowers with elevated floral humidity (figure 1). Temperature differences between flowers were negligibly small but could also not account for this finding. This is because in the passive artificial flower preference test, should temperature differences be directing foraging decisions, we would have expected to find the reverse of the preferences observed (Dyer et al., 2006), as dry passive flowers were slightly warmer than humid variants. Our finding aligns well with previous observations in the hawkmoth *H. lineataIa* (von Arx et al., 2012) and field observations of alpine fly pollinators in India (Nordström et al., 2017) which suggested preferences for flowers with higher floral humidity.

When rewarding and nonrewarding flowers differed in humidity cues bumblebees showed greatly enhanced foraging success with experience compared to bees in the control group (figure 2), where rewarding and nonrewarding flowers did not differ in humidity. This indicates floral humidity differences enhanced learning of rewarding flowers. Bumblebees showed an initial preference to more humid flowers but with experience learned to favour visits to the rewarding flower type, whether rewarding flowers were producing higher or lower levels of floral humidity. In the latter case, bees were trained against their initial preference, nevertheless their performance did not differ much from the group rewarded on the higher humidity flowers, which suggests that it is not difficult for bees to learn to favour less humid flowers if they are more rewarding.

Our results indicate that floral humidity represents a floral signal or cue that can be used far more widely than previously thought. The hygrosensitive ceolocapitular sensilla of generalist pollinators like bees are likely to be able to respond to both the amount of humidity produced by the flower itself (Yokohari, 1983; Yokohari et al., 1982) and the rate of change in humidity experienced as the bee approaches or passes the flowers (Tichy and Kallina, 2014, 2010). This ability to distinguish different levels of flower humidity has important consequences, given that natural flowers can differ very strongly in floral humidity. The presence, absence and difference in floral humidity between flowers are likely to be an integral part of the multisensory displays and transmit valuable information that bumblebees can respond to and learn whilst detecting, choosing or handling flowers. Consequently, traits that influence the floral humidity production may be adaptive to plants (Harrap et al., 2020a). Traits that increase floral humidity levels may increase visitation by naïve bumblebees, positively influencing their unlearned preferences and, by creating differences in humidity between flowers, aid the learning and recognition of flowers from those that produce less humidity. Nevertheless, floral traits that suppress floral humidity production may still exist in natural flowers, as these traits can also be adaptive. Although naïve bees may be less attracted to flowers with reduced floral humidity, the absence of humidity can be easily learned by pollinators since it represents a predictive cue for floral rewards, as we show here. For example, previous work has shown that *Vinca herbacea* and *Linum grandiflorum* showed no floral humidity or less humidity than extraneous humidity sources (Harrap et al., 2020a). Based on the findings of our learning experiments we would predict that bumblebees detect such lack of humidity and would very easily distinguish these flowers from humidity-producing species. Similar adaptations of the floral display that go against naïve bee preferences but may enhance floral recognition have been observed previously in non-blue coloured flowers (Dyer and Chittka, 2004; Gumbert et al., 1999; Lynn et al., 2005) and cold flowers (Whitney et al., 2008).

Our findings extend the understanding of plant-pollinator interaction but also shed a light on a novel function of humidity perception in insects. As an environmental cue for insects (Abou-Shaara et al., 2017; Enjin, 2017), humidity can have important influences on levels of foraging activity (McCall and Primack, 1992; Peat and Goulson, 2005), (micro)habitat selection (particularly when avoiding desiccation) (Enjin, 2017; Knecht et al., 2017; Perttunen and Erkkilä, 1952; Sun et al., 2018), selection of oviposition sites (Okal et al., 2013), locating vertebrate hosts (Chappuis et al., 2013; Olanga et al., 2010; Smart and Brown, 1956), nest maintenance (Human et al., 2006; Nicolson, 2009), and context-dependent flight responses (Wolfin et al., 2018). We show here that this well-developed sensory capacity can also be used in a flower-foraging context when processing and learning multimodal information that is available in flower displays. Like many insects (Liu et al., 2007), other bees (Fialho et al., 2014) possess hygrosensitive receptors. However, bumblebees in particular have a more widespread distribution of hygrosensative receptors across their antennae than other bees (Fialho et al., 2014) and this may result in differences in sensitivity that might determine the extent to which different pollinators can make use of floral humidity cues.

Due to floral humidity having strong links to nectar evaporation, floral humidity has been proposed as an ‘honest signal’ (von Arx, 2013; von Arx et al., 2012). In *O. caespitosa* flowers, removal or blocking of floral nectar reduced the intensity of floral humidity. Honest signals correspond with the reward state of flowers, indicating temporary rewardlessness to pollinators (*e*.*g*. due to a recent visitation by a pollinator), allowing pollinators to avoid wasteful visits to unrewarding flowers, increasing pollinator efficiency and preference to honest signallers (Knauer and Schiestl, 2015; von Arx, 2013). The spontaneous and learned responses to floral humidity demonstrated here may allow bumblebees to adjust visitation to favour rewarding flowers. However, it is uncertain whether floral humidity intensity directly indicates flower reward state, and therefore functions honestly, in all species that produce it.

The multisensory displays of most flowers are dominated by visual and olfactory cues. Within multimodal floral displays, humidity preferences might have complementary effects rather than directly guiding decisions and responses. The spontaneous floral humidity preferences of naïve bumblebees were subtle despite the differences in humidity between the artificial flowers being comparable to larger differences observed between natural flowers (Harrap et al., 2020a). Humidity differences generally may therefore enhance the capacity of pollinators to learn multimodal floral stimuli, independently how reliably the presence of nectar can be inferred. It has been shown that learning of one signal can be improved when other signal modalities are present, even when they do not differ with that initial signal (Clarke et al., 2013; Kunze and Gumbert, 2001; Leonard et al., 2011a). This may be due to contextualising other signals or changing perceptual saliences when presented together (Leonard et al., 2011b; Raguso, 2004). For naïve pollinators, humidity cues could be particularly useful as it would help to localise flowers more accurately, which could scaffold their learning and improve their foraging success (Raine and Chittka, 2008). Now that more widespread pollinators, bumblebees, have demonstrated that capacity to respond to biologically relevant levels of floral humidity, further investigation of how floral humidity is used as part of a multimodal display will provide a more holistic understanding of plant-pollinator interactions in nature.

## Acknowledgements

We are grateful to Alanna Kelly for technical assistance in the laboratory.

## Data, Code and Materials

Raw data, data plotted in figures as well as the annotated *R* code and excel object files (where appropriate) necessary to generate graphical figures and repeat analysis are available in the supplementary materials. The datafiles pertaining to measurement and analysis of artificial flower humidity are found in supplementary file 2. The datafiles pertaining bumblebee behavioural experiments are found in the supplementary file 3. Within both these .zip files there is a word document that provides a key to datasheets and R files contained within.

## Authors’ contributions

HMW, NHdI and SAR acquired funding for research, conceived of project ideas, and supervised research. HDK and MJMH collected artificial flower humidity data. MJMH designed methodology, collected bee behavioural data and carried out analysis. MJMH and SAR lead writing of the manuscript. All authors contributed to manuscript drafts and gave final approval for manuscript submission.

