## Supplementary file 1 for "Bumblebees can detect floral humidity"

**SUPPLEMENTARY INFORMATION**

**SUPPLEMENTARY INFORMATION APPENDIX 1: Artificial flower design and maintenance**

*Artificial flower design: active flowers*

Active flowers were similar in design to those utilised by von Arx et al. (2012) but modified to better suit bumblebee foragers. These artificial flowers had a flower top with small holes (figure S1A) to a chamber below the flower head (figure S1B). This chamber was connected by 6mm external, 4mm internal diameter airline tubing (*MARINA blue airline*, Hagen, Mansfield, USA) to a pump assembly outside the foraging arena (figure S1D and S1E). In humidity-producing flowers, airflow was through water in a bubbler in this pump assembly that elevated humidity of the air that was fed to the flower head. Less-humid dry flowers were also created, where the pump assembly was the same but the bubbler was empty. Thus, airflow at the flower head was the same between flower variants but the air reaching the flower head in dry flowers had not had its humidity increased.

A full schematic of an active artificial flower’s pump assembly and its installation in the flight arena is given in figure S1E. Airflow from a mechanical fish tank air pump (*MARINA cool 11135*, Hagen, Mansfield, USA) was connected to a bubbler chamber by a 22cm section airline. The last 7cm of this 22cm tube was inserted within the bubbler chamber and the last 2cm of this section of tube was cut away at a 20º angle. This allowed the tube from the pump to be below the water level and allowed surface tension at the end of the pipe to be weaker promoting movement of bubbles through water when the bubbler was full.

This bubbler chamber was made with an airtight 150ml tupperware cylinder (made with either ‘*Snac-Pacs food tubes*’, Wilko, Worksop, UK or ‘*Snack tubes*’, Smash Nude Food Movers, Mitcham, Australia). Two holes were drilled into the lid of this and fitted with rubber grommets to match tube diameters. This chamber would be filled with either 100ml of water (that had been allowed to settle at room temperature overnight) in humid flower variants, or left empty in dry flower variants. This meant that in humidity producing flowers air that had undergone mixing with the water travelled up to the top of the bubbler, while flow of air continued in the same way in dry flowers but without humidity being increased in this air.

A 26cm section of airline tubing was then connected to a rotameter (*Omega FL-3802C*, *Omega engineering*, Manchester, UK). Only enough tubing of this section to clear the grommet was inserted into the bubbler chamber (3mm). This meant that this tube was always above the water level and collected humid air (and dryer air) collecting at the top of the bubbler. These rotameters regulate airflow using a screw to obstruct airflow. Airflow was set at 2.69ml s^-1^, controlling the flow of humid or dry air to the artificial flower head. The rotameter output was linked to a 90cm long section of tubing that entered the flight arena through holes in a wooden bracket installed on the doorways of the foraging arenas. This 90cm tube would link to the artificial flower itself. Eight active flowers were used at any one time. Four would enter the arena from either side through two door brackets (figure S1D).

A 25mm diameter hole would be cut into the bottom of a plastic cup (Dart C71-130, Huntingdon, UK), and a 6mm diameter hole was punched 3mm from the top on one side, just above the lip on the top. This cup was upturned, and all but the lip was covered with black electrical tape (figure S1C). This functioned as the flower stand, holding the artificial flowers upright. The 90cm tube from the rotameter was fed into the 6mm hole and up through the 25mm hole. This base was then weighted using modelling clay allowing it to stand in place against any elastic tension created by bending the tube.

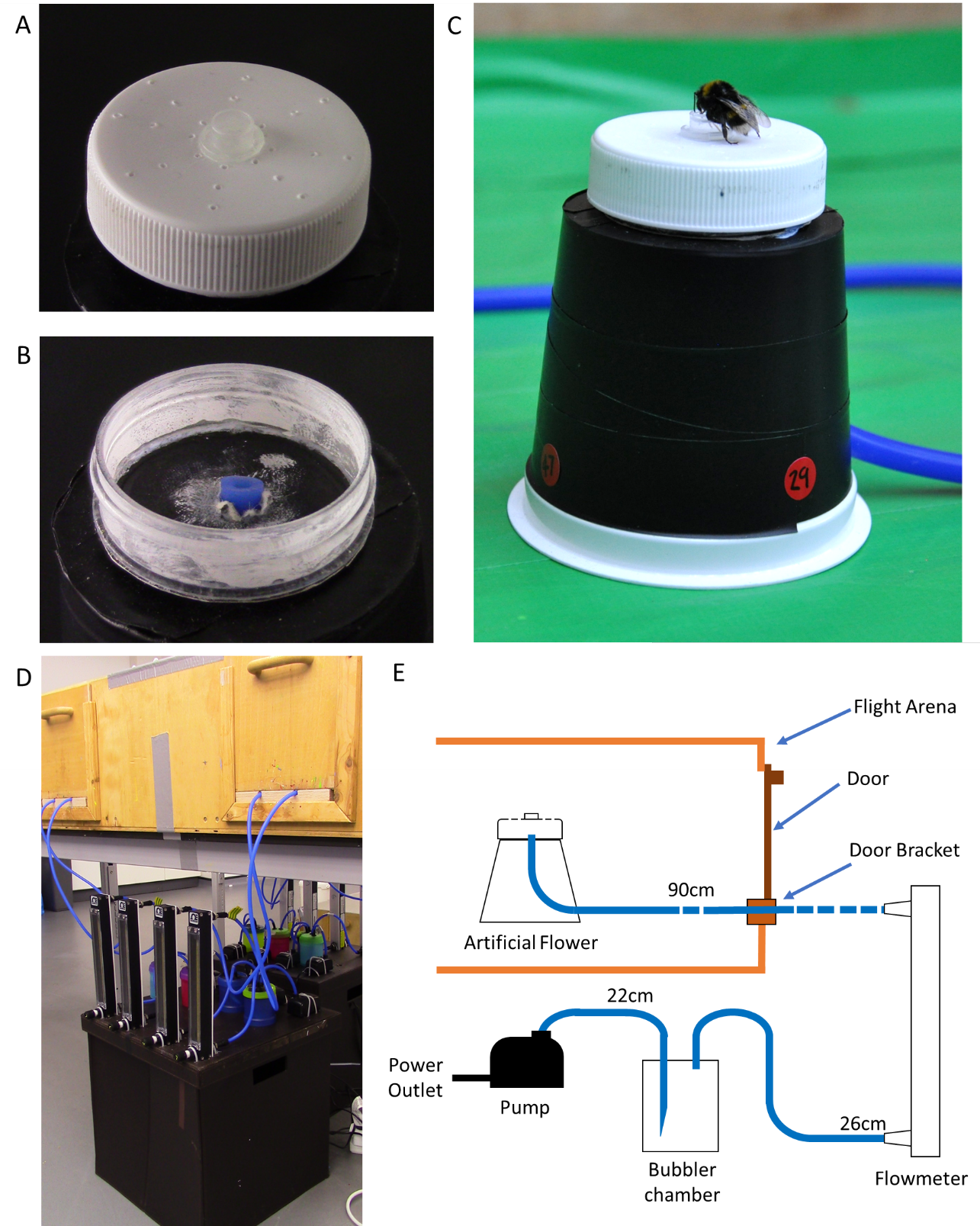

**Figure S1:** The active artificial flowers used in bumblebee experiments. Panel **A**) The artificial flower head. Note the holes on the flower head for air to escape. **B**) The flower head with the head unscrewed, allowing the chamber under and pipe entry point to be seen. **C**) Bumblebee feeding from active humidity flower as they appear in the flight arena. **D**) The pump-bubbler-rotameter assembly installed below the flight arena. Note the rubber tubes entering the arena through brackets below the doors. **E**) A diagrammatical representation of each artificial flower and its pump mechanism and how it installs through the flight arena through a door bracket. Rubber tubes are represented by blue lines connecting components, the lengths of tubes are given above each tube.

The head of the artificial flower was made from a specimen jar (Thermo scientific sterilin, PS 60ml, with white plastic lids), where the top 1cm of the jar (containing the screw lid thread) was cut away. A 0.5ml Eppendorf tube lid (Hamburg, Germany) was upturned and stuck down in the centre of the jar top, to function as the feeding well containing sucrose solution or water. 24 holes were made in the jar lid using a thumbtack pin. These 24 holes were in lines of 3, each line being at a 45^o^ angle from the next, the first hole at the base of the feeding well, the others separated by 5mm (figure S1A). The screw thread was stuck to thick card using super glue (Precision super glue, Loctite, Hemel Hempstead, UK). Once dry, the joint of this card and the screw thread was covered in glue in order to ensure as good a seal as possible. A 6mm diameter hole was then punched through the centre of this card base, and the last 3mm of the 90cm tube leading to the rotameter was inserted through it (figure S1B) and secured with electrical tape. The flower lid was then screwed tight and the tubing pulled taut so that the flower head would rest on the stand. A small amount of *blue tac* was stuck to the underside of the flower head, to hold it in place against the stand. Artificial flowers thus appeared to bees as the jar lid on top of a trapezoid base (figure S1C).

To aid identification of individual flowers by experimenters, in a way bees would not be able to identify, red sticky dots were stuck about the base of the flower stand, and two-digit numbers written on these with black permanent marker (figure S1C). These numbers were odd on half the flowers entering the foraging arena on each side, even on the other half of the flowers (*i.e*. two of each side’s four flowers were even, two were odd). The black on red colours of these numbers would be hard for bumblebees to make out given their visual systems (Davies et al., 2013). Additionally, these numbers were two digit, this allowed the initial digit to be even number in odd number stickers and *vice versa*. This meant bees were unlikely to recognise flower based on the number shapes (if they can be seen at all) as even and odd digits were present on all flowers. As the bubblers that contained water could be changed, whether even or odd numbered flowers corresponded with humid or dry flower variants could be changed between experiment days.

*Artificial flower design: passive flowers*

Passive flowers created humidity by evaporation of water from components internal to the flower through a permeable lid. In dry, less humid artificial flower variants, construction was identical but without water being added to the flowers internal components.

Passive artificial flowers were built from a specimen jar (Thermo scientific sterilin, PS 60ml, with white plastic lids). The bodies of the jars were covered with black electrical tape in the same way described in the ‘small artificial flowers’ in Harrap et al. (2017), to prevent bees visually identifying the artificial flowers by contents. Flowers were numbered with randomly generated numbers, as described in Harrap et al. (2017), to allow visual identification of humid or dry variant flowers by human experimenters (figure S2). Again these odd and even numbers had several digits, including even and odd digits. This reduced the chance of bees identifying rewards based on the shape of numbers, as they occurred on both even and odd numbered flowers.

A 35 mm circular hole was cut into the centre of each jar’s lid, and the edges were smoothed to remove any excess plastic. This hole removed most of the flat top of the jar but maintained the screw threading assembly of the jar lid (figure S2B and S2C). The top surfaces of the artificial flowers were made with a sheet of fine gauze material (made from cut out segments of TERESIA curtains, IKEA, Leiden, Netherlands) stretched over the jar aperture and screwed into place using the cut away lid screw assembly. Any excess gauze visible below the screw lid on all flowers was cut away. This created a gauze top to the artificial flower slightly lower than the plastic rim of the artificial flower (figure S2D). This gauze surface was firm enough for the bee to walk upon, would help obscure jar contents, and was permeable to the evaporation produced by internal components of artificial flowers (see below). An upturned 0.5ml eppendorf tube lid was painted black and placed in the centre of the gauze indentation, functioning as a feeding well during experiments. This lid was not stuck down and could be moved by the bees while feeding, however it was too heavy for the bees to easily lift and the plastic rim of the artificial flower prevented bees upturning the lid or pushing it off the artificial flower (figure S2E).

Three discs of 1cm thick sponge were placed inside the specimen jars within each artificial flower. These discs (cut from cellulose sponge wipes, Co-op, Manchester, UK) were 40 mm diameter, the width of the specimen jar, (figure S2A). The top (visible) sponges were all identical green. For humid artificial flower these discs were wetted prior to experiments, and at the midpoint of conditioning experiments, as per the protocol laid out in the following section. The evaporation from this wet sponge increased the relative humidity above these artificial flowers. Dry artificial flowers did not have any water added to the sponges.

Each batch of 24 sponge discs were stored in a sealed bag following being cut from sheets until needed. As each flower needed 3 discs and 8 flowers were presented to the bee during trials (see main text), all the discs used in one day were from the same sponge batch and stored in the same way. All sponge discs were discarded after a day of use.

*Artificial flower setup*

Before preference experiments using active artificial flowers, the pump assembly for four artificial flowers was placed under the flight arena table on both the right and left sides of the arena (as shown for one side in figure S1D). This allowed artificial flowers to be placed in the arena through door brackets placed in the doors on that side. On each side two of the artificial flowers had odd numbering, two even numbering (making eight flowers in total, four odd, four even). The bubbler chambers of either odd or even numbered flowers were filled with 100ml of water, the other dry, as described above. In order to ensure a good seal on the Tupperware and the input and output for the rotameter and grommet seals for the bubbler all these seals were strengthened with electrical tape. The airflow on all rotameters was then set to a 2.69ml s^-1^ using the rotameter screw seal.

Passive artificial flowers were prepared as follows before each bee’s trial in both preference and learning experiments. Sponge discs for dry artificial flower variants were inserted as they were from the bag into the specimen jar before the gauze and flower lid were screwed in. Sponge discs for humid artificial flower variants were submerged until sodden in a pitcher of water, that had been allowed to settle at the lab’s temperature overnight, before insertion into the specimen jar and screwing down of gauze and lid. To avoid conflating indicators of which flowers contained wet sponge this was done so that the gauze remained dry. If the gauze got wet at any part of the experiment it was removed and replaced with a fresh dry sheet. Once sponge discs were inserted and tops screwed on, the humidity produced by humid flowers was checked using a handheld hygrometer (Maplin Electronics, UK). If the relative humidity 5mm above the artificial flowers did not read at least 2% higher than the ambient humidity of the lab using this hygrometer, sponge discs would be removed and re-soaked. As humid flowers contained water, inserting of sponge and evaporation may cause a drop in temperature, artificial flower temperature was checked before trials began using a thermal camera (FLIR systems, Inc., Wilsonville, USA). During all thermal imaging emissivity parameter value used were 0.95 (Infrared Training Center, 2008) and lab reflected temperature was measured using a tin foil mirror (Harrap et al., 2018) to have a consistent value of 20ºC. As the water used for sponge wetting had been allowed to settle at room temperature, humid flower variants and dry flowers rarely differed in temperature enough for bees to detect (where detectability is presumed to occur if the temperature difference is more than 2ºC (Heran, 1952)). However, if the humid flowers differed in temperature from the dry flowers by more than 1ºC, whichever flower variant was warmer would be cooled by placing them on a tray inside a refrigerator at 5ºC until the temperature difference between flower variants was below 1ºC. If both these humidity and temperature requirements were met flowers were ready to be presented to bees and experiments could start. During our learning experiments passive artificial flowers were also be re-prepared (as described above) at the end of the foraging bout when the bee crossed the halfway point in terms of visit number (35 visits or more). A foraging bout constituting the time between a bee leaving the nest to forage in the flight arena and exiting the arena to return to the nest.

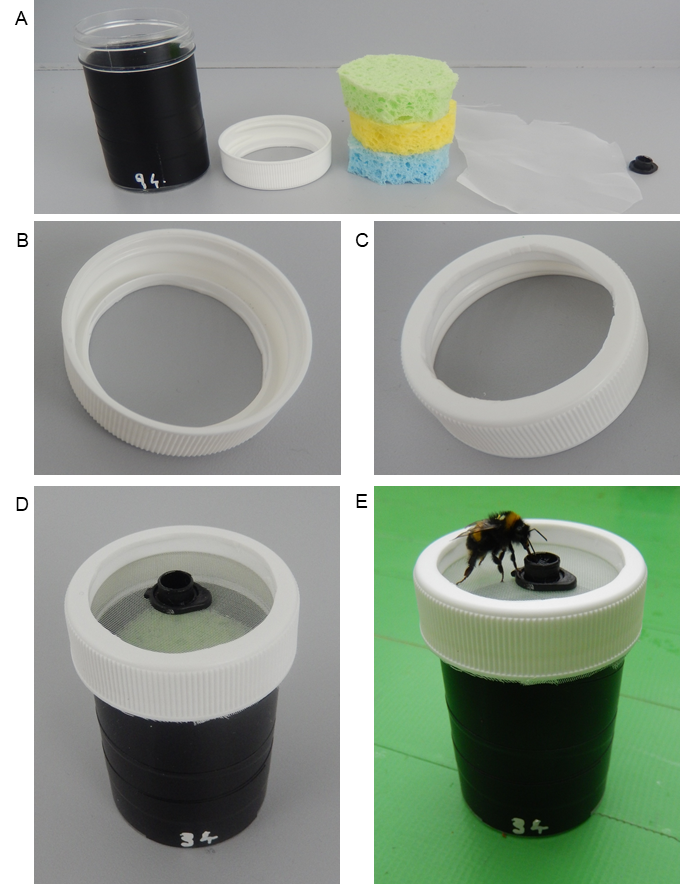

**Figure S2:** The passive artificial flowers used in bumblebee experiments. **A**) The artificial flower components, from left to right: the specimen jar; the specimen jar lid; three sponge discs; gauze fabric; and an Eppendorf tube lid. **B**) The specimen jar lid showing the cut away section, leaving the screw assembly, and the lip, from below. **C**) The same, but from above. **D**) The assembled artificial flower. **E**) Bumblebee feeding from the artificial flowers.

*Artificial flower cleaning and maintenance*

Both artificial flower types were cleaned regularly throughout the experiments to prevent any conflating scent marks left behind be bee visits (Pearce et al., 2017; Stout and Goulson, 2001). Cleaning occurred at the end of each foraging bout.

When active artificial flowers were cleaned, all flowers were removed from the arena and the tops were wiped with ethanol, with care taken to not apply liquid over the holes. The tubing prevented flowers from being moved to completely different locations during tests, due to tubes being linked to door brackets, so following cleaning the door bracket holes each artificial flower entered the arena by were changed on each side (*i.e*. a single flower would now enter the arena from a different hole on the same side of the arena). As tupperware seals, the tightness of the rotameter screw, rotameter input or output seals and grommet seals can weaken under the pressure system of the pump assembly, rotameter airflow was checked after any flower cleaning and rotameter adjusted. Where necessary other seals were repaired to maintain a 2.69ml s^-1^ airflow rate.

When passive artificial flowers were cleaned there was a risk that the fabric top of the flower retained scent better than plastic parts. Furthermore, returning a passive artificial flower to the arena with an ethanol wetted top may conflate the humidity differences between flowers under consideration in foraging tests. Consequently, when flowers were removed from the arena for cleaning, the lids and gauze were removed. The plastic parts of the lip were wiped down with ethanol, and a fresh sheet of unused gauze was screwed down onto the flower top with the clean lid. Excess gauze outside of the screw assembly would be cut away as before (figure S2D). This cleaning and replacement of fabric prevented scent marks that might aid reward discrimination from accumulating on the flower tops and allowed artificial flowers to remain consistently dry to the touch of the bees between foraging bouts.

**SUPPLEMENTARY INFORMATION APPENDIX 2: Measurement and Evaluation of Artificial Flower Floral Humidity**

*Sampling artificial floral humidity*

Both humidity-producing and dry variants of both artificial flower types, eight of each active artificial flower variant and twelve of each passive artificial flower variant, were sampled using the robot arm transect based method described in Harrap et al. (2020a) to evaluate the artificial floral humidity they produce.

This method utilizes a *Staubli* RX 160 robot arm (Pfäffikon, Switzerland). This robot arm carries out paired transects of the flower headspace of (upward facing) flowers placed on a table in front of it (see (Harrap et al., 2020) for detailed schematics of the robot sampling area setup and transects). First, an ‘*x* axis transect’ sampling humidity across the horizontal span of flower. Second, a ‘*z* axis transect’ sampling humidity vertically upwards from the flower. Along these transects the robot samples relative humidity using a DHT-22 humidity probe (Aosong Electronics, Huangpu, China) mounted upon it. Sampling positions of these transects are resolved autonomously by the robot relative to a manually input transect central point. This transect central point being a space 5mm above the flower centre. Simultaneously to humidity sampling by the robot, the background humidity of the sampling area is measured by a background humidity probe on the table flowers are placed upon. The robot carries out these paired transects on each flower presented to it in turn, carrying out a probe calibration step after sampling each flower. In this probe calibration step the robot moves its humidity probe to the same position as the background humidity probe. Here the two probes sample the same location, and thus sample an area of equal humidity. Measurements taken in this probe calibration step are used to account for differences in reading between probes (±5% according to the manufacturer’s specifications), allowing this source of error to be removed (see (Harrap et al., 2020)). The robot then repeats this sequence, sampling each flower in turn (with probe calibrations) three more times.

Four artificial flowers (two humid, two dry variants of the same artificial flower type) would be sampled each day. This means each artificial flower’s headspace was sampled four times over approximately 21 hours within roughly 308 minute intervals. Feeding wells of artificial flowers were filled with a 25μl droplet of water at the start of robot arm sampling, as bees would normally encounter flowers with water or sucrose solution present in the feeding well (see main text). Thus, it was necessary to understand what humidity the flowers produce with this water present. All artificial flowers show upwards orientation, therefore no reorientation of the flowers (see (Harrap et al., 2020)) was necessary. Flowers were placed on the table for sampling. The pump assembly of active artificial flowers was set up under the table within the sampling area of the robot. Flowers were otherwise set up as described in Supplementary Information Appendix 1. Passive humid flowers were not re-wetted at any time after initial setup during floral humidity sampling by the robot.

From these transects we calculated change in humidity relative to the background humidity ($\Delta RH$) across the transects. Once the robot has stopped at a measurement point on each transect the arm measures humidity c.100 times in 200 seconds. These c.100 measurements taken at each measurement point have been found to have high repeatability of each other (Harrap et al., 2020). Therefore, the mean $\Delta RH$ of each measurement point along the transects was used for analysis. Linear models that allowed differing humidity structures and changes in humidity with replicate transects, were fitted to the $\Delta RH$ data of each artificial flower variant, as done in (Harrap et al., 2020) for different flower species. Models of different humidity structure that allowed humidity structure and/or intensity to change with replicate transects or not were fitted to the x and z axis $\Delta RH$ transect data. Models fitted to the x axis data allowed either a quadratic, linear or flat relationship, depending on the model, between transect position in the x axis and $\Delta RH$. Models fitted to the z axis data allowed a logarithmic or flat relationship between transect position in the z axis and $\Delta RH$. Throughout all models artificial flower identity was included as a random factor influencing floral humidity intensity. Further details of the models fit to humidity data can be seen in the code attached to the datafiles of this manuscript. Best fitting models for $\Delta RH$ across the *x* and *z* axis humidity transects of each artificial flower variant were found using AIC. Summary values $X_{t}^{max}$ (the point in the *x* axis transect where humidity difference is greatest) and ${\Delta RH}_{x}^{max}$ (the greatest mean humidity difference generated) according to the best fitting models of each artificial flower variant were calculated. Further detail of summary value calculation, can be found in (Harrap et al., 2020).

*Assessment of artificial flower humidity*

The best fitting models, $X_{t}^{max}$ and ${\Delta RH}_{x}^{max}$ values for humid and dry variants of each flower type are given in Supplementary Information table S1. Additionally, the parameter values for the best fitting models of humid and dry variants of each flower type are given in Supplementary Information table S2.

Humidity produced by the humid variants of both active (figure S3) and passive flowers (figure S4) was similar in intensity to floral humidity produced by real flower species with ${\Delta RH}_{x}^{max}$ values greater than 3% (table S1), such as *Eschscholzia californica* (3.24%), *Taraxacum* agg. (3.35%), or *Ranunculus acris* (3.41%). Some humidity came from water droplets in the well of the artificial flowers, explaining how humidity was still produced by dry flower variants. The humidity produced by the feeding well was likely to be lower in dry active flowers than the dry passive flowers due to the effect of the airflow in active flowers dispersing water vapour (figures S3 and S4). However, this production of smaller amounts of humidity in the dry variants of both active and passive artificial flowers was both lower than humid variants, and similar to that produced by real flowers (Harrap et al., 2020). Active dry flowers produced humidity differences similar to *Convolvulus sabatius* (0.87%), *Cyanus segetum* (1.10%) and *Linum usitatissium* (0.8%)*.* Passive Dry flowers produced humidity in the range of flowers producing moderate amounts of floral humidity such as *Leucanthemum vulgare* (1.79%) and *Achillea millefolium* (1.73%). The differences in relative humidity intensity between humid and dry flower variants was 2.16% in active flowers and 1.36% in passive flowers. These differences in humidity intensity were similar to those observed between different flowers (Harrap et al., 2020), which means that the floral humidity levels that the experimental bees were exposed to were within the bounds of differences they might experience when foraging on natural flowers.

Spatially, the humidity was distributed in similar ways to that found in flowers (Harrap et al., 2020). Peak values were measured in the central area when probing in the horizontal plane across the upward-facing surface (*x* axis, figure S3A and B, figure S4A and B), and declined when moving away upwards from the surface (*z* axis, figure S3C and D, figure S4C and D).

The humidity produced by both variants of active flowers remained largely stable throughout the *circa* 21 hour sampling period, during which the humidity of their headspace was sampled four times (figure S3). The only change observed during this period was a drop in humidity of the dry active flowers after the initial transect (figure S3A), which was affected more strongly by the evaporation of the water in the feeding well during the initial transect. As dry active flowers were regularly refilled throughout bee experiments after being emptied (see main text), it is likely that the humidity differences were maintained at levels shown in the initial transect. So, the mean difference in humidity intensity (in terms of ${\Delta RH}_{x}^{max}$) between dry and humid active flower variants remained ~2.16% during experiments. The passive flowers were less stable, with the floral humidity regularly dropping with replicate transects in the dry flower variant (figure S4A) and dropping after the second replicate transect and again after the third in the humid flower variant (figure S4B). This was caused by the drying out of wet sponge components as well as the evaporation of water from the feeding well. During the experiments with bees the feeding wells of passive flowers were refilled and, where appropriate, sponge components re-wetted (see main text and supplementary information appendix 1). As the passive humid flowers show stable average humidity intensities for the first and second transect replicates, this means that the initial peak in humidity lasted for at least ten hours before drying out affects humidity intensity. Preference and learning trials rarely took this long, so it is unlikely that the humidity would drop much below the initial intensities in the time allowed between re-wetting. Thus, the mean difference in humidity intensity (in terms of ${\Delta RH}_{x}^{max}$) between dry and humid passive flower variants remained at ~1.36% within the timescales of our experiments.

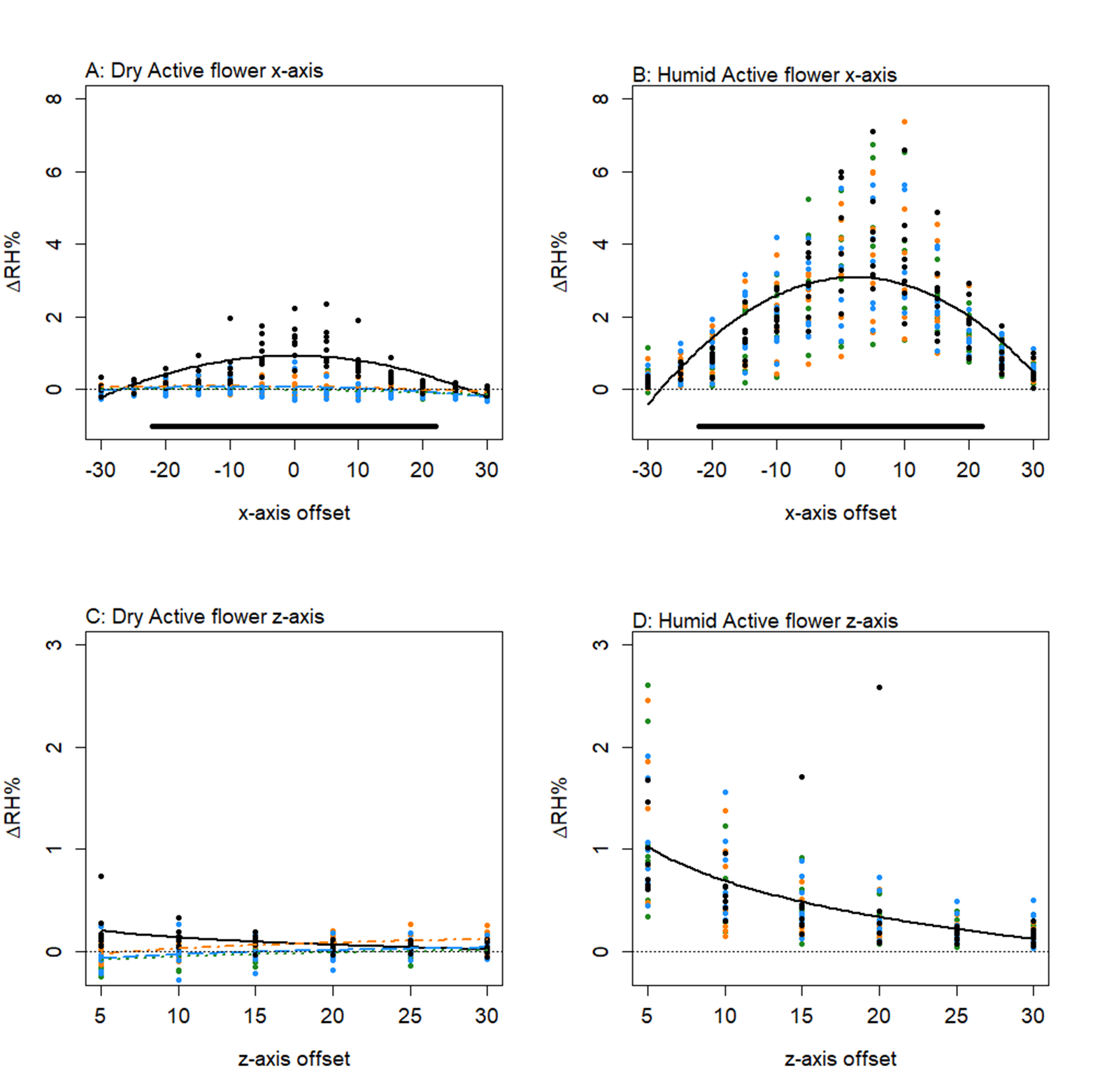
**Figure S3:** The difference in humidity relative to the background ($\Delta RH$) for transects of active flowers. *x* and *z* axis transects are given for: dry active flower variant in **A** and **C** respectively; and for the humid active flower variant in **B** and **D** respectively. All axis offsets are relative to the transect central point and in millimetres. The thin dotted line indicates a 0% change in humidity (the background level). Bold lines indicate the mean change in humidity as predicted by the best fitting model for that flower. Colour and dashing of bold lines and points indicate the replicate transect: solid black, first transect; long-dash blue, second transect; dash-dot orange, third transect; dotted green, fourth transect. The solid bar above the *x* axis transects indicate the diameter of the flower top (44mm) relative to the *x* axis. Number of active flowers of each variant sampled, *n* = 8.

**Figure S4:** The difference in humidity relative to the background ($\Delta RH$) for transects of passive flowers. *x* and *z* axis transects are given for: dry passive flower variant in **A** and **C** respectively; and for the humid passive flower variant in **B** and **D** respectively. See figure S3 for description of figure details. Number of passive flowers of each variant sampled, *n* = 12.

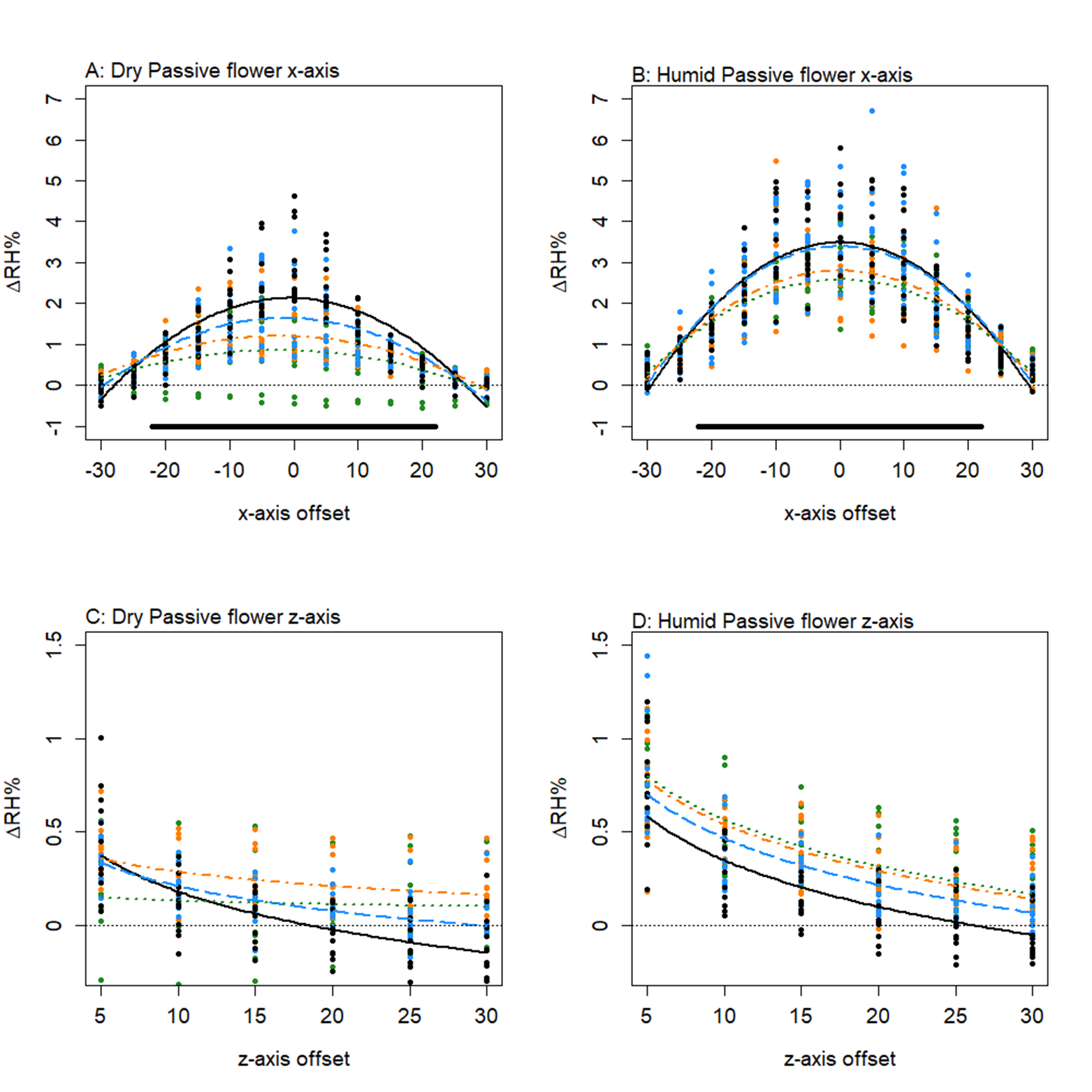

**Table S1:** The best fitting models, $X_{t}^{max}$ and $\Delta{RH}_{x}^{max}$ values for the both variants of both types of artificial flowers. Subscript letters following best fitting models indicate shape of that model: L, linear models; and Q, quadratic models (note quadratic models were not fitted to the z axis). Subscript values next to $x_{t}^{max}$ values indicate replicate effects: the number itself referring to the replicate transect at which ${\Delta RH}_{x}^{max}$ was found; no subscript values indicates replicate transects have no effect on humidity. For further details on models, see (Harrap et al., 2020) and the code attached to the manuscript.

| **Artificial Flower Type** | **Flower variant** | **Best fitting model for the** | | $\boldsymbol{X}_{\boldsymbol{t}}^{\boldsymbol{max}}$ | $\boldsymbol{\Delta RH}_{\boldsymbol{x}}^{\boldsymbol{max}}$ |
| --- | --- | --- | --- | --- | --- |
|  |  | ***x* axis** | ***z* axis** |  |  |
| Active | Humid | m3_Q_ | z1_L_ | 2.19 | 3.08 |
|  | Dry | m10_Q_ | z4_L_ | -0.12_1_ | 0.92 |
| Passive | Humid | m9_Q_ | z3_L_ | 0_1_ | 3.49 |
|  | Dry | m10_Q_ | z4_L_ | -0.61_1_ | 2.13 |

**Table S2:** The parameter values of the best fitting models of both *x* and *z* axis models from our analysis of humidity structure of each variant of each artificial flower type. Parameters are identified by the R model fixed effect labels, column ‘R’, and the parameter names given for the equivalent parameters in (Harrap et al., 2020), column ID, to facilitate comparison. For further detail on parameters of models and parameter function consult Harrap et al (2020) and the code attached to the manuscript. All values are given in scientific format (*g*·E*x* = *g*·10*^x^*).

| **Flower type** | | **Active flowers** | | **Passive flowers** | |
| --- | --- | --- | --- | --- | --- |
| **Flower Variant** | | **Humid** | **Dry** | **Humid** | **Dry** |
| *ID* | *R* |  |  |  |  |
| *I_x_* | (intercept) | 3.07 E+00 | 9.25 E-01 | 3.49 E+00 | 2.13 E+00 |
| *A_x_* | *xoffset* | 1.48 E-02 | -2.96 E-04 |  | -3.50 E-03 |
| *B_x_* | *x2* | -3.38 E-03 | -1.26 E-03 | -4.05 E-03 | -2.85 E-03 |
| *r*_2_*_x_* | *rep1* |  | -8.52 E-01 | -1.01 E-01 | -4.89 E-01 |
| *r*_3_*_x_* | *rep2* |  | -8.52 E-01 | -6.91 E-01 | -9.35 E-01 |
| *r*_4_*_x_* | *rep3* |  | -9.34 E-01 | -9.11 E-01 | -1.28 E+00 |
| *g*_2_*_x_* | *xoffset:rep1* |  | -2.64 E-03 |  | -2.24 E-03 |
| *g*_3_*_x_* | *xoffset:rep2* |  | -1.98 E-03 |  | -2.26 E-03 |
| *g*_4_*_x_* | *xoffset:rep3* |  | -2.23 E-03 |  | -1.40 E-03 |
| *c*_2_*_x_* | *x2:rep1* |  | 1.06 E-03 | 3.17 E-04 | 7.83 E-04 |
| *c*_3_*_x_* | *x2:rep2* |  | 1.18 E-03 | 1.16 E-03 | 1.60 E-03 |
| *c*_4_*_x_* | *x2:rep3* |  | 1.20 E-03 | 1.51 E-03 | 1.92 E-03 |
| *I_z_* | (intercept) | 2.01 E+00 | 4.04 E-01 | 1.27 E+00 | 9.39 E-01 |
| *B_z_* | *lnzoffset* | -5.50 E-01 | -1.12 E-01 | -3.86 E-01 | -3.16 E-01 |
| *r*_2_*_z_* | *rep1* |  | -5.87 E-01 | 1.17 E-01 | -2.32 E-01 |
| *r*_3_*_z_* | *rep2* |  | -5.89 E-01 | 1.93 E-01 | -3.68 E-01 |
| *r*_4_*_z_* | *rep3* |  | -5.96 E-01 | 2.17 E-01 | -7.36 E-01 |
| *c*_2_*_z_* | *lnzoffset:rep1* |  | 1.76 E-01 |  | 1.09 E-01 |
| *c*_3_*_z_* | *lnzoffset:rep2* |  | 2.01 E-01 |  | 1.97 E-01 |
| *c*_4_*_z_* | *lnzoffset:rep3* |  | 1.71 E-01 |  | 2.87 E-01 |

**SUPPLEMENTARY INFORMATION APPENDIX 3: Statistical models and simplification procedure for bee learning experiments**

The following represents our full model used for analysis of bee learning before any simplification was applied, these models are identical to those employed in (Harrap et al., 2017, p. 2018) with parameter identities modified to suit the test groups used in the current study:

$y_{nx}=i+\left( \ln x*l \right)+D\left( s_{d}+\left( \ln x*c_{d} \right) \right)+H\left( s_{h}+\left( \ln x*c_{h} \right) \right)+\left( b_{n}+\left( \ln x*r_{n} \right) \right)$.	(S.1)

Where $y_{nx}$ is the arcsine square root success rate of bee $n$ over the previous 10 visits to the artificial flowers, at $x$ flower visits. $x$ is the number of the visits the bee has made to the artificial flowers, the data for $y$ is calculated in blocks of 10 visits (i.e. at 10, 20, 30, 40, 50, 60 and 70 visits). Parameter $i$is the initial arcsine square root success rate, the intercept, for bees in the control group when $x=0$. Parameter $l$ dictates the change in arcsine square root success rate with increased $x$ in the control group, thus $l$ is effectively the learning speed parameter and allows bee’s experience to effect success rate. $D$ and $H$ are Boolean parameters which allow the model to alter $y$ depending on which test group the bee is in. $D$ indicates whether the bee is in the dry rewards test group, where:

$D=\left\{ \begin{aligned} 0 \\ 1 \end{aligned}\begin{matrix} bee is not in the dry rewards group, \\ bee is in the dry rewards group, \end{matrix} \right.$	(S.2)

$H$ indicates whether the bee is in the humid rewards test group, where:

$H=\left\{ \begin{aligned} 0 \\ 1 \end{aligned}\begin{matrix} bee is not in the humid rewards group, \\ bee is in the humid rewards group. \end{matrix} \right.$	(S.3)

$s_{d}$ and $s_{h}$ are the change in initial arcsine square root success rate, relative to $i$, for bees in the dry and humid rewards test groups respectively. $c_{d}$ and $c_{h}$ are the change in learning speed, relative to $l$, for bees in the dry and humid rewards test groups respectively. Variation between individual bees was included in our model as a random factor. $b_{n}$ and $r_{n}$ represent the change in initial arcsine square root success rate and learning speed, for bee number $n$. In the model described in equation S.1 parameters $i, l, s_{d}, s_{h}, c_{d}, c_{h}, b_{n}$ and $r_{n}$ are parameters to be estimated.

Model simplification procedure involved paired comparisons between the standing ‘best model’, beginning with the full model described in equation S.1, with a simpler model. Simpler models were constructed from the standing best model but with further parameters removed (effectively forcing the relevant parameters to equal zero) in the order described below. Should the simpler model have a lower AIC or be comparable to the standing best fitting models based on AIC, as laid out by (Richards, 2008), this simpler model would become the best model for the next comparison. If removal of a parameter led to a significant increase in AIC, again as laid out by (Richards, 2008), the standing best (more complex) model would remain the best for the next comparison.

Initially the effects of random factors were compared, a model without $r_{n}$ was compared to the complete model. This allowed testing of whether individual bees differed only in intercepts or intercepts and learning speed (as in the full model). In both experiments $r_{n}$ had no significant effect on the model, and is thus not included in subsequent models below. Secondly interaction effects were investigated by removing $c_{d}$ and $c_{h}$. This created a model where the shape of the relationship between $x$ and $y$ in all test groups was dictated only by $l$.

Should the best model, according to AIC, find no significant interaction the effects of the test groups would be investigated by removing $D$ and $H$ creating a model were all test group both showed the same intercepts and learning. Finally, the impact of experience on success was compared by removing the learning parameter $l$. Should the best fitting interaction model include interaction effects individual models for each test group would be fitted as follows:

$y_{nx}=i+\left( \ln x*l \right)+b_{n}$.	(S.4)

For each test group, using the model described in equation S.4, we tested whether bee foraging success changed with the number of visits the bee has made by removing parameter $l$.

An annotated copy of the code used in this analysis is available attached to the datafiles associated with this manuscript.

**SUPPLEMENTARY INFORMATION APPENDIX 4: AIC tables and sampling dates of artificial flower floral humidity analyses**

For each individual artificial flower of each variant the date and time at which the first x axis transect replicate began is given (YYYY-MM-DD-hh-mm-ss). In each AIC table, each species having one for x and z axis models, AIC and degrees of freedom ‘df’ are given: see (Harrap et al., 2020) for description of the different models. Difference in ΔAIC, here calculated as AIC of model with the lowest AIC minus that of the current model, is also provided. Within each AIC table, shaded and in bold are the best fitting models as per the guidelines given in (Richards, 2008).

| **Active Humid** | | | |  |  | | | |
| --- | --- | --- | --- | --- | --- | --- | --- | --- |
| X axis model | df | AIC | ΔAIC |  | Z axis model | df | AIC | ΔAIC |
| **m3** | **5** | **1114.88** | **0.00** |  | **z1** | **4** | **86.67** | **0.00** |
| m7 | 8 | 1119.73 | -4.85 |  | z3 | 7 | 92.09 | -5.42 |
| m10 | 14 | 1127.27 | -12.39 |  | z4 | 10 | 95.24 | -8.57 |
| m2 | 4 | 1151.32 | -36.44 |  | z0 | 3 | 233.29 | -146.62 |
| m6 | 7 | 1156.27 | -41.39 |  | z2 | 6 | 239.03 | -152.36 |
| m9 | 10 | 1158.71 | -43.82 |  |  |  |  |  |
| m1 | 4 | 1473.78 | -358.90 |  | Sampling dates | | 2017-11-16-11-08-52 | |
| m5 | 7 | 1479.30 | -364.42 |  |  | | 2017-11-16-12-26-15 | |
| m8 | 10 | 1485.08 | -370.20 |  |  |  | 2017-11-16-13-43-38 | |
| m0 | 3 | 1488.09 | -373.21 |  |  |  | 2017-11-16-15-01-05 | |
| m4 | 6 | 1493.63 | -378.75 |  |  |  | 2018-02-01-11-54-27 | |
|  |  |  |  |  |  |  | 2018-02-01-10-37-04 | |
|  |  |  |  |  |  |  | 2018-02-01-13-11-52 | |
|  |  |  |  |  |  |  | 2018-02-01-14-29-17 | |
| **Active Dry** | | | |  |  | | | |
| X axis model | df | AIC | ΔAIC |  | Z axis model | df | AIC | ΔAIC |
| **m10** | **14** | **-29.10** | **0.00** |  | **z4** | **10** | **-347.76** | **0.00** |
| m9 | 10 | -22.24 | -6.85 |  | z3 | 7 | -312.58 | -35.18 |
| m7 | 8 | 126.84 | -155.94 |  | z2 | 6 | -310.97 | -36.78 |
| m6 | 7 | 132.75 | -161.84 |  | z1 | 4 | -279.22 | -68.53 |
| m5 | 7 | 205.87 | -234.96 |  | z0 | 3 | -278.31 | -69.44 |
| m8 | 10 | 210.19 | -239.28 |  |  |  |  |  |
| m4 | 6 | 210.35 | -239.45 |  | Sampling dates | | 2017-11-20-15-10-07 | |
| m3 | 5 | 314.15 | -343.25 |  |  | | 2017-11-20-16-27-30 | |
| m2 | 4 | 317.09 | -346.18 |  |  |  | 2017-11-20-17-44-55 | |
| m1 | 4 | 364.45 | -393.55 |  |  |  | 2017-11-20-19-02-19 | |
| m0 | 3 | 366.80 | -395.90 |  |  |  | 2018-01-30-10-35-56 | |
|  |  |  |  |  |  |  | 2018-01-30-11-53-23 | |
|  |  |  |  |  |  |  | 2018-01-30-13-10-46 | |
|  |  |  |  |  |  |  | 2018-01-30-14-28-06 | |

| **Passive Humid** | | | |  |  | | | |
| --- | --- | --- | --- | --- | --- | --- | --- | --- |
| X axis model | df | AIC | ΔAIC |  | Z axis model | df | AIC | ΔAIC |
| **m9** | **10** | **1208.27** | **0.00** |  | z4 | 10 | -125.11 | 0.00 |
| m10 | 14 | 1214.90 | -6.62 |  | **z3** | **7** | **-122.95** | **-2.15** |
| m6 | 7 | 1260.35 | -52.08 |  | z1 | 4 | -76.13 | -48.98 |
| m7 | 8 | 1261.82 | -53.54 |  | z2 | 6 | 111.63 | -236.74 |
| m2 | 4 | 1298.20 | -89.93 |  | z0 | 3 | 129.54 | -254.64 |
| m3 | 5 | 1299.70 | -91.43 |  |  |  |  |  |
| m4 | 6 | 2047.90 | -839.63 |  | Sampling dates | | 2017-10-03-10-34-17 | |
| m5 | 7 | 2049.75 | -841.48 |  |  | | 2017-10-03-13-09-02 | |
| m0 | 3 | 2054.29 | -846.01 |  |  |  | 2017-10-04-10-58-13 | |
| m8 | 10 | 2055.56 | -847.28 |  |  |  | 2017-10-04-12-15-36 | |
| m1 | 4 | 2056.14 | -847.87 |  |  |  | 2017-10-05-11-36-06 | |
|  |  |  |  |  |  | | 2017-10-05-14-10-52 | |
|  |  |  |  |  |  |  | 2017-10-10-13-23-13 | |
|  |  |  |  |  |  | | 2017-10-10-14-40-36 | |
|  |  |  |  |  |  |  | 2017-10-23-12-10-47 | |
|  |  |  |  |  |  |  | 2017-10-23-13-28-10 | |
|  |  |  |  |  |  |  | 2017-10-24-10-17-19 | |
|  |  |  |  |  |  |  | 2017-10-24-12-52-05 | |
| **Passive Dry** | | | |  |  | | | |
| X axis model | df | AIC | ΔAIC |  | Z axis model | df | AIC | ΔAIC |
| **m10** | **14** | **872.95** | **0.00** |  | **z4** | **10** | **-156.34** | **0.00** |
| m9 | 10 | 890.60 | -17.65 |  | z3 | 7 | -131.26 | -25.08 |
| m7 | 8 | 1001.48 | -128.54 |  | z1 | 4 | -103.48 | -52.86 |
| m6 | 7 | 1019.29 | -146.34 |  | z2 | 6 | -67.79 | -88.56 |
| m3 | 5 | 1102.00 | -229.05 |  | z0 | 3 | -46.80 | -109.54 |
| m2 | 4 | 1116.68 | -243.74 |  |  |  |  |  |
| m5 | 7 | 1468.42 | -595.48 |  | Sampling dates | | 2017-10-03-11-51-40 | |
| m8 | 10 | 1474.10 | -601.16 |  |  |  | 2017-10-03-14-26-27 | |
| m4 | 6 | 1475.71 | -602.76 |  |  |  | 2017-10-04-13-32-59 | |
| m1 | 4 | 1514.24 | -641.29 |  |  | | 2017-10-04-14-50-22 | |
| m0 | 3 | 1520.78 | -647.83 |  |  |  | 2017-10-05-10-18-43 | |
|  |  |  |  |  |  | | 2017-10-05-12-53-29 | |
|  |  |  |  |  |  |  | 2017-10-10-10-48-25 | |
|  |  |  |  |  |  |  | 2017-10-10-12-05-48 | |
|  |  |  |  |  |  |  | 2017-10-23-10-53-22 | |
|  |  |  |  |  |  | | 2017-10-23-14-45-33 | |
|  |  |  |  |  |  |  | 2017-10-24-11-34-42 | |
|  |  |  |  |  |  |  | 2017-10-24-14-09-28 | |
