## Supplementary file 2 for "Bumblebees can detect floral humidity": Artificial Floral Humidity Data guide.docx

Attached is the floral humidity data collected by the robot arm transects described in the main text. For each artificial flower type (active or passive) and variant (dry or humid) two csv files are provided. The ‘Raw’ datafile, indicated by ‘[*artificial flower type*] [*flower variant*] Raw’ contains all individual humidity measurements taken throughout sampling of all transects pertaining to each artificial flower kind and variant (i.e. all the individual measurements made in each measurement period). The ‘Means’ datafile, indicated by ‘[*artificial flower type*] [*flower variant*] Means’ contains the mean averages of each measurement period pertaining to each artificial flower type and variant. These mean values are used in assessments of floral humidity as described in the main text and are the data plotted in the floral humidity graphs in figures 1 and 2.

Finally a ‘.R’ code file ‘Artificial flower code.R’ is provided. Within this R file is sufficient code for the analysis of floral humidity reported, and the code needed to generate Figure S3 and S4’s panels. Annotation within the R file is present to guide users through it.

**Guide to Column headings: ‘Raw’ datafile**

| [unlabeled] | - | A counter of humidity measurement datapoints within the file. |
| --- | --- | --- |
| X. | - | Humidity measurement number. This number is applied by the humidity probes and counts up with each measurement but is reset periodically. |
| Species | - | The artificial flower type and variant of the measurement. |
| Individual | - | The individual flower identification number within each few sampling days – This value resets across each few days of sampling, but is used to generate the totIndividual values that identify individual flowers across all humidity sampling days. |
| totIndividual | - | The individual flower identification number across all humidity sampling - $n$ within model equations. |
| date | - | The time and date of the measurement, formatted as *yyyy-mm-dd-hh-mm-ss.* |
| sample.. | - | The replicate transect number of the measurement point. 0 = first transect, 1 = second transect, 2 = third transect, 3 = forth transect. |
| Sample.order | - | The number of the flower individual within the randomly selected sequence in which flowers were sampled. 0 = first flower in sampling order, 1 = second flower in sampling order, 2 = third flower in sampling order, 3 = forth flower in sampling order. |
| x.or.z.transect | - | The indicator of the transect being carried out– x or z |
| xoffset | - | The x axis offset of the current measurement. |
| zoffset | - | The z axis offset of the current measurement. |
| Background.Humidity | - | The background humidity measurement. |
| Background.Temperature | - | The background temperature measurement - The temperature measurement taken by the background probe. |
| Focal.Humidity | - | The uncorrected focal humidity measurement |
| Focal.Temperature | - | The temperature measurement taken by the focal probe. |
| Corrected.Focal.humidity | - | The corrected focal humidity measurement. |
| Change.in.RH. | - | The change in humidity between focal and background probes ($\Delta RH$) |
| block | - | The current measurement period number – this value counts up across all sampling. |

**Guide to Column headings: ‘Means’ datafile**

| [unlabeled] | - | A counter of datapoints within the file. |
| --- | --- | --- |
| Block | - | The measurement period |
| Sp. | - | The artificial flower type and variant of the measurement. |
| xORz | - | Indicator as to transect being carried out – x or z |
| Individual | - | The individual flower identification number within each few sampling days – This value resets across each few days of sampling, but is used to generate the totIndividual values that identify individual flowers across all humidity sampling days. |
| totIndividual | - | The individual flower identification number across all humidity sampling - $n$ within model equations. |
| sample.. | - | The replicate transect number of the measurement point. 0 = first transect, 1 = second transect, 2 = third transect, 3 = forth transect. |
| Sample.order | - | The number of the flower individual within the randomly selected sequence in which flowers were sampled. 0 = first flower in sampling order, 1 = second flower in sampling order, 2 = third flower in sampling order, 3 = forth flower in sampling order. |
| xoffset | - | The x axis offset of the current measurement period. |
| zoffset | - | The z axis offset of the current measurement period. |
| x2 | - | The x axis offset squared (used in analyses). |
| z2 | - | The z axis offset squared (ultimately not used in any analyses). |
| Background.Humidity | - | The mean background humidity measurement of the measurement period. |
| Background.Temperature | - | The mean background temperature measurement of the measurement period. |
| Focal.Humidity | - | The mean uncorrected focal humidity measurement of the measurement period. |
| Focal.Temperature | - | The mean temperature measurement taken by the focal probe of the measurement period. |
| Change.in.RH. | - | The mean change in humidity between focal and background probes ($\Delta RH$) of the measurement period. |
