## Supplementary file 2 for "Bumblebees can detect floral humidity": Artificial flower code.R

```
#Artificial flower humidity analysis code from-
#Bumblebees can detect floral humidity

#Authors:
#Michael J. M. Harrap
#Natalie Hempel de Ibarra
#Henry D. Knowles
#Heather M. Whitney
#Sean A. Rands

#Below is the code for the analysis of the bee preference data for the
#aforementioned publication. Instruction and explanation of code is given in
# sections marked with hashtag (#). Outputs from preceding code are given in
#section marked with inverted commas ('').

#For further explanation of the data and analysis see the above publication.

#All analysis was carried out in R version 3.6.3 (2020-02-29) -- "Holding the Windsock"

# clear memory and start afresh
rm(list = ls())

#set working directory to file 'Artificial flower humidity data' file
setwd(".../Artifical flower humidity data") #replace '...' with your directory

#Load packages
library(lme4)

#Load datafiles
PassiveDrymeans<-read.csv(file = "Passive Dry Means.csv", header = TRUE)
PassiveHumidmeans<-read.csv(file = "Passive Humid Means.csv", header = TRUE)

ActiveDrymeans<-read.csv(file = "Active Dry Means.csv", header = TRUE)
ActiveHumidmeans<-read.csv(file = "Active Humid Means.csv", header = TRUE)

#The below analysis is carried out on a datasheet named Focalmeans,
#Which artificial flower type and variant is analysed is assigned using
#the below code

#SET THE FLOWER VARIENT AND TYPE FOR ANALYSIS
Focalmeans<-ActiveHumidmeans
#where the artificial flower tpye and varient to be analysed can be inserted to the right of '<-'
#i.e. it can be any of the follwoing:
#'PassiveDrymeans' 'PassiveHumidmeans' 'ActiveDrymeans' 'ActiveHumidmeans'
  
#following assignation of the Focalmeans it is then necessary to run the following code to
#separate data of different transects, transform z offset and confirm rep is assigned as a factor.

FocalmeansREP0<-subset(Focalmeans, subset = sample..=="0")
FocalmeansREP0X<-subset(FocalmeansREP0, subset =xORz==" x")
FocalmeansREP0Z<-subset(FocalmeansREP0, subset =xORz==" z")

FocalmeansREP1<-subset(Focalmeans, subset = sample..=="1")
FocalmeansREP1X<-subset(FocalmeansREP1, subset =xORz==" x")
FocalmeansREP1Z<-subset(FocalmeansREP1, subset =xORz==" z")

FocalmeansREP2<-subset(Focalmeans, subset = sample..=="2")
FocalmeansREP2X<-subset(FocalmeansREP2, subset =xORz==" x")
FocalmeansREP2Z<-subset(FocalmeansREP2, subset =xORz==" z")

FocalmeansREP3<-subset(Focalmeans, subset = sample..=="3")
FocalmeansREP3X<-subset(FocalmeansREP3, subset =xORz==" x")
FocalmeansREP3Z<-subset(FocalmeansREP3, subset =xORz==" z")


FocalX<-rbind(FocalmeansREP0X,FocalmeansREP1X,FocalmeansREP2X,FocalmeansREP3X)
Focalz<-rbind(FocalmeansREP0Z,FocalmeansREP1Z,FocalmeansREP2Z,FocalmeansREP3Z)
Focal<-rbind(FocalmeansREP0,FocalmeansREP1,FocalmeansREP2,FocalmeansREP3)

Focal$lnzoffset<-log(Focal$zoffset+1)
FocalX$lnzoffset<-log(FocalX$zoffset+1)
Focalz$lnzoffset<-log(Focalz$zoffset+1)

Focal$rep<-factor(Focal$sample..)
FocalX$rep<-factor(FocalX$sample..)
Focalz$rep<-factor(Focalz$sample..)

is.factor(Focal$rep)
is.factor(FocalX$rep)
is.factor(Focalz$rep)


#Once the above step is carried out for a artifical flower type and vatient 
#the below models can be run to fit models to the flower type and varient assinged to focalmeans.
#model content and strucutre are described in references given in the main text.

#Xaxis models
m0<-lmer(Change.in.RH.~(1|totIndividual),data=FocalX, REML = FALSE)

m1<-lmer(Change.in.RH.~xoffset+(1|totIndividual),data=FocalX, REML = FALSE)

m2<-lmer(Change.in.RH.~x2+(1|totIndividual),data=FocalX, REML = FALSE)
m3<-lmer(Change.in.RH.~xoffset+x2+(1|totIndividual),data=FocalX, REML = FALSE)

m4<-lmer(Change.in.RH.~rep+(1|totIndividual),data=FocalX, REML = FALSE)
m5<-lmer(Change.in.RH.~xoffset+rep+(1|totIndividual),data=FocalX, REML = FALSE)
m6<-lmer(Change.in.RH.~x2+rep+(1|totIndividual),data=FocalX, REML = FALSE)
m7<-lmer(Change.in.RH.~xoffset+x2+rep+(1|totIndividual),data=FocalX, REML = FALSE)

m8<-lmer(Change.in.RH.~xoffset+rep+rep:xoffset+(1|totIndividual),data=FocalX, REML = FALSE)
m9<-lmer(Change.in.RH.~x2+rep+rep:x2+(1|totIndividual),data=FocalX, REML = FALSE)
m10<-lmer(Change.in.RH.~xoffset+x2+rep+rep:xoffset+rep:x2+(1|totIndividual),data=FocalX, REML = FALSE)

AIC(m0,m1,m2,m3,m4,m5,m6,m7,m8,m9,m10)

#models can be viewed in detail using
#summary(MODEL NAME HERE)
summary(m3)

#AIC Output for passive dry artifical flowers
'    df       AIC
m0   3 1520.7752
m1   4 1514.2387
m2   4 1116.6836
m3   5 1101.9983
m4   6 1475.7072
m5   7 1468.4223
m6   7 1019.2885
m7   8 1001.4820
m8  10 1474.1016
m9  10  890.5952
m10 14  872.9454'

#AIC Output for passive humid artifical flowers
'    df      AIC
m0   3 2054.286
m1   4 2056.141
m2   4 1298.200
m3   5 1299.701
m4   6 2047.901
m5   7 2049.753
m6   7 1260.353
m7   8 1261.817
m8  10 2055.556
m9  10 1208.274
m10 14 1214.895'


#AIC Output for active dry artifical flowers
'    df       AIC
m0   3 366.79928
m1   4 364.45288
m2   4 317.08619
m3   5 314.14889
m4   6 210.35445
m5   7 205.86540
m6   7 132.74524
m7   8 126.84445
m8  10 210.18539
m9  10 -22.24340
m10 14 -29.09649'


#AIC Output for active humid artifical flowers
'    df      AIC
m0   3 1488.090
m1   4 1473.777
m2   4 1151.320
m3   5 1114.882
m4   6 1493.632
m5   7 1479.301
m6   7 1156.270
m7   8 1119.728
m8  10 1485.083
m9  10 1158.706
m10 14 1127.274'

#

#zAXIS MODEL

z0<-lmer(Change.in.RH.~(1|totIndividual),data=Focalz, REML = FALSE)

z1<-lmer(Change.in.RH.~lnzoffset+(1|totIndividual),data=Focalz, REML = FALSE)

z2<-lmer(Change.in.RH.~rep+(1|totIndividual),data=Focalz, REML = FALSE)
z3<-lmer(Change.in.RH.~lnzoffset+rep+(1|totIndividual),data=Focalz, REML = FALSE)

z4<-lmer(Change.in.RH.~lnzoffset+rep+rep:lnzoffset+(1|totIndividual),data=Focalz, REML = FALSE)

AIC(z0,z1,z2,z3,z4)


#models can be viewed in detail using
#summary(MODEL NAME HERE)
summary(z1)


#AIC Output for passive dry artifical flowers
'   df        AIC
z0  3  -46.80232
z1  4 -103.48256
z2  6  -67.78591
z3  7 -131.26456
z4 10 -156.34285'

#AIC Output for passive humid artifical flowers
'   df        AIC
z0  3  129.53896
z1  4  -76.12792
z2  6  111.63493
z3  7 -122.95437
z4 10 -125.10597'

#AIC Output for active dry artifical flowers
'   df       AIC
z0  3 -278.3103
z1  4 -279.2233
z2  6 -310.9741
z3  7 -312.5750
z4 10 -347.7552'

#AIC Output for active humid artifical flowers
'   df       AIC
z0  3 233.29369
z1  4  86.67054
z2  6 239.03474
z3  7  92.08927
z4 10  95.23924'


#________________________________________________________________________


#Code for graphs
rm(list = ls())

#set working directory to file 'Artifical flower humidity data' file
setwd("C:/Users/mjmha/Desktop/ProcB/Artifical flower humidity data")

#read datafiles,
Drymeans<-read.csv(file = "Passive Dry Means.csv", header = TRUE)
Wetmeans<-read.csv(file = "Passive Humid Means.csv", header = TRUE)

Flow10Dry<-read.csv(file = "Active Dry Means.csv", header = TRUE)
Flow10Wet<-read.csv(file = "Active Humid Means.csv", header = TRUE)


#Passive Dry
#subset data by axis and replicate transect
DrymeansREP0<-subset(Drymeans, subset = sample..=="0")
DrymeansREP0X<-subset(DrymeansREP0, subset =xORz==" x")
DrymeansREP0Z<-subset(DrymeansREP0, subset =xORz==" z")

DrymeansREP1<-subset(Drymeans, subset = sample..=="1")
DrymeansREP1X<-subset(DrymeansREP1, subset =xORz==" x")
DrymeansREP1Z<-subset(DrymeansREP1, subset =xORz==" z")

DrymeansREP2<-subset(Drymeans, subset = sample..=="2")
DrymeansREP2X<-subset(DrymeansREP2, subset =xORz==" x")
DrymeansREP2Z<-subset(DrymeansREP2, subset =xORz==" z")

DrymeansREP3<-subset(Drymeans, subset = sample..=="3")
DrymeansREP3X<-subset(DrymeansREP3, subset =xORz==" x")
DrymeansREP3Z<-subset(DrymeansREP3, subset =xORz==" z")

#X
#set model lines
Rep0dry<-curve(2.1294919+(x*-0.0034951)+((x^2)*-0.0028499),xlim=c(-30,30),ylim=c(-8,8))
Rep1dry<-curve(2.1294919+ -0.4890497+(x*(-0.0034951+-0.0022388))+((x^2)*((-0.0028499+0.0007830))),xlim=c(-30,30),ylim=c(-8,8))
Rep2dry<-curve(2.1294919+-0.9351942+(x*(-0.0034951+-0.0022643))+((x^2)*((-0.0028499+0.0016043))),xlim=c(-30,30),ylim=c(-8,8))
Rep3dry<-curve(2.1294919+-1.2828812+(x*(-0.0034951+-0.0014013))+((x^2)*((-0.0028499+0.0019170))),xlim=c(-30,30),ylim=c(-8,8))

#Figure S4A
plot(Change.in.RH.~xoffset,data=DrymeansREP3X,xlab="x-axis offset",ylab=expression(paste("",Delta,"RH%")), pch=20,col="forestgreen",ylim=c(-1,7),yaxt="n",cex.axis=1.2,cex.lab=1.2)
axis(2,at=c(-2,-1,0,1,2,3,4,5,6,7,8),labels=c("-2","-1","0","1","2","3","4","5","6","7","8"),cex.axis=1.2)

mtext(expression(paste("A: Dry Passive flower x-axis")),adj=0,cex=1.2)

points(Change.in.RH.~xoffset,data=DrymeansREP2X,pch=20,col="darkorange1")
points(Change.in.RH.~xoffset,data=DrymeansREP1X,pch=20,col="dodgerblue")
points(Change.in.RH.~xoffset,data=DrymeansREP0X,pch=20,col="black")

lines(Rep0dry,lwd=2, lty=1)
lines(Rep1dry,col="dodgerblue",lwd=2, lty=5)
lines(Rep2dry,col='darkorange1',lwd=2, lty=4)
lines(Rep3dry,col='forestgreen',lwd=2, lty=3)
abline(0,0,lty=3)
segments(-22.,-1,22.,-1,lwd=5,col="black")


#Z
#set model lines
Rep0zdry<-curve(0.93932+((log(x+1))*-0.31643),xlim=c(5,30),ylim=c(0,4))
Rep1zdry<-curve(0.93932-0.23189+((log(x+1))*(-0.31643+0.10887)),xlim=c(5,30),ylim=c(0,4))
Rep2zdry<-curve(0.93932-0.36820+((log(x+1))*(-0.31643+0.19726)),xlim=c(5,30),ylim=c(0,4))
Rep3zdry<-curve(0.93932-0.73589+((log(x+1))*(-0.31643+0.28652)),xlim=c(5,30),ylim=c(0,4))

#Figure s4C
plot(Change.in.RH.~zoffset,data=DrymeansREP3Z,xlab="z-axis offset",ylab=expression(paste("",Delta,"RH%")), pch=20,col="forestgreen",ylim=c(-0.25,1.5),yaxt="n",cex.axis=1.2,cex.lab=1.2)
axis(2,at=c(0,0.5,1,1.5,2),labels=c("0","0.5","1","1.5","2"),cex.axis=1.2)

mtext(expression(paste("C: Dry Passive flower z-axis")),adj=0,cex=1.2)

points(Change.in.RH.~zoffset,data=DrymeansREP2Z,pch=20,col="darkorange1")
points(Change.in.RH.~zoffset,data=DrymeansREP1Z,pch=20,col="dodgerblue")
points(Change.in.RH.~zoffset,data=DrymeansREP0Z,pch=20,col="black")

lines(Rep0zdry,lwd=2, lty=1)
lines(Rep1zdry,col="dodgerblue",lwd=2, lty=5)
lines(Rep2zdry,col="darkorange1",lwd=2, lty=4)
lines(Rep3zdry,col="forestgreen",lwd=2, lty=3)
abline(0,0,lty=3)

#Passive Humid
#subset data by axis and replicate transect
WetmeansREP0<-subset(Wetmeans, subset = sample..=="0")
WetmeansREP0X<-subset(WetmeansREP0, subset =xORz==" x")
WetmeansREP0Z<-subset(WetmeansREP0, subset =xORz==" z")

WetmeansREP1<-subset(Wetmeans, subset = sample..=="1")
WetmeansREP1X<-subset(WetmeansREP1, subset =xORz==" x")
WetmeansREP1Z<-subset(WetmeansREP1, subset =xORz==" z")

WetmeansREP2<-subset(Wetmeans, subset = sample..=="2")
WetmeansREP2X<-subset(WetmeansREP2, subset =xORz==" x")
WetmeansREP2Z<-subset(WetmeansREP2, subset =xORz==" z")

WetmeansREP3<-subset(Wetmeans, subset = sample..=="3")
WetmeansREP3X<-subset(WetmeansREP3, subset =xORz==" x")
WetmeansREP3Z<-subset(WetmeansREP3, subset =xORz==" z")

#X
#set model lines
Rep0wet<-curve(3.4949653+(x*0)+((x^2)*-0.0040525),xlim=c(-30,30),ylim=c(-8,8))
Rep1wet<-curve(3.4949653+-0.1013831+(x*(0))+((x^2)*((-0.0040525+0.0003167))),xlim=c(-30,30),ylim=c(-8,8))
Rep2wet<-curve(3.4949653+-0.6906035+(x*(0))+((x^2)*((-0.0040525+0.0011635))),xlim=c(-30,30),ylim=c(-8,8))
Rep3wet<-curve(3.4949653+-0.9107727+(x*(0))+((x^2)*((-0.0040525+0.0015120))),xlim=c(-30,30),ylim=c(-8,8))

#Figure S4b
plot(Change.in.RH.~xoffset,data=WetmeansREP3X,xlab="x-axis offset",ylab=expression(paste("",Delta,"RH%")), pch=20,col="forestgreen",ylim=c(-1,7),yaxt="n",cex.axis=1.2,cex.lab=1.2)
axis(2,at=c(-2,-1,0,1,2,3,4,5,6,7,8),labels=c("-2","-1","0","1","2","3","4","5","6","7","8"),cex.axis=1.2)

mtext(expression(paste("B: Humid Passive flower x-axis")),adj=0,cex=1.2)

points(Change.in.RH.~xoffset,data=WetmeansREP2X,pch=20,col="darkorange1")
points(Change.in.RH.~xoffset,data=WetmeansREP1X,pch=20,col="dodgerblue")
points(Change.in.RH.~xoffset,data=WetmeansREP0X,pch=20,col="black")

lines(Rep0wet,lwd=2, lty=1)
lines(Rep1wet,col="dodgerblue",lwd=2, lty=5)
lines(Rep2wet,col='darkorange1',lwd=2, lty=4)
lines(Rep3wet,col='forestgreen',lwd=2, lty=3)
abline(0,0,lty=3)
segments(-22.,-1,22.,-1,lwd=5,col="black")


#Z
#set model lines
Rep0zwet<-curve(1.27482+((log(x+1))*-0.38631),xlim=c(5,30),ylim=c(0,4))
Rep1zwet<-curve(1.27482+0.11725+((log(x+1))*(-0.38631)),xlim=c(5,30),ylim=c(0,4))
Rep2zwet<-curve(1.27482+0.19285+((log(x+1))*(-0.38631)),xlim=c(5,30),ylim=c(0,4))
Rep3zwet<-curve(1.27482+0.21745+((log(x+1))*(-0.38631)),xlim=c(5,30),ylim=c(0,4))

#Figure s4d
plot(Change.in.RH.~zoffset,data=WetmeansREP3Z,xlab="z-axis offset",ylab=expression(paste("",Delta,"RH%")), pch=20,col="forestgreen",ylim=c(-0.25,1.5),yaxt="n",cex.axis=1.2,cex.lab=1.2)
axis(2,at=c(0,0.5,1,1.5,2),labels=c("0","0.5","1","1.5","2"),cex.axis=1.2)

mtext(expression(paste("D: Humid Passive flower z-axis")),adj=0,cex=1.2)

points(Change.in.RH.~zoffset,data=WetmeansREP2Z,pch=20,col="darkorange1")
points(Change.in.RH.~zoffset,data=WetmeansREP1Z,pch=20,col="dodgerblue")
points(Change.in.RH.~zoffset,data=WetmeansREP0Z,pch=20,col="black")

lines(Rep0zwet,lwd=2, lty=1)
lines(Rep1zwet,col="dodgerblue",lwd=2, lty=5)
lines(Rep2zwet,col="darkorange1",lwd=2, lty=4)
lines(Rep3zwet,col="forestgreen",lwd=2, lty=3)
abline(0,0,lty=3)


#Dry active flowers
#
#subset data by axis and replicate transect
Flow10DrymeansREP0<-subset(Flow10Dry, subset = sample..=="0")
Flow10DrymeansREP0X<-subset(Flow10DrymeansREP0, subset =xORz==" x")
Flow10DrymeansREP0Z<-subset(Flow10DrymeansREP0, subset =xORz==" z")

Flow10DrymeansREP1<-subset(Flow10Dry, subset = sample..=="1")
Flow10DrymeansREP1X<-subset(Flow10DrymeansREP1, subset =xORz==" x")
Flow10DrymeansREP1Z<-subset(Flow10DrymeansREP1, subset =xORz==" z")

Flow10DrymeansREP2<-subset(Flow10Dry, subset = sample..=="2")
Flow10DrymeansREP2X<-subset(Flow10DrymeansREP2, subset =xORz==" x")
Flow10DrymeansREP2Z<-subset(Flow10DrymeansREP2, subset =xORz==" z")

Flow10DrymeansREP3<-subset(Flow10Dry, subset = sample..=="3")
Flow10DrymeansREP3X<-subset(Flow10DrymeansREP3, subset =xORz==" x")
Flow10DrymeansREP3Z<-subset(Flow10DrymeansREP3, subset =xORz==" z")


#X
#Generate model lines
Rep0dry10<-curve(9.246e-01+(x*-2.955e-04)+((x^2)*-1.264e-03),xlim=c(-30,30),ylim=c(-8,8))
Rep1dry10<-curve(9.246e-01+-8.519e-01+(x*(-2.955e-04 +-2.640e-03))+((x^2)*((-1.264e-03+1.058e-03))),xlim=c(-30,30),ylim=c(-8,8))
Rep2dry10<-curve(9.246e-01+-8.520e-01+(x*(-2.955e-04 +-1.977e-03))+((x^2)*((-1.264e-03+1.179e-03))),xlim=c(-30,30),ylim=c(-8,8))
Rep3dry10<-curve(9.246e-01+-9.340e-01+(x*(-2.955e-04 +-2.225e-03))+((x^2)*((-1.264e-03+1.198e-03))),xlim=c(-30,30),ylim=c(-8,8))

#Figure s3A
plot(Change.in.RH.~xoffset,data=Flow10DrymeansREP3X,xlab="x-axis offset",ylab=expression(paste("",Delta,"RH%")), pch=20,col="forestgreen",ylim=c(-1,8),yaxt="n",cex.axis=1.2,cex.lab=1.2)
axis(2,at=c(-2,0,2,4,6,8),labels=c("-2","0","2","4","6","8"),cex.axis=1.2)

points(Change.in.RH.~xoffset,data=Flow10DrymeansREP2X,pch=20,col="darkorange1")
points(Change.in.RH.~xoffset,data=Flow10DrymeansREP1X,pch=20,col="dodgerblue")
points(Change.in.RH.~xoffset,data=Flow10DrymeansREP0X,pch=20,col="black")

mtext(expression(paste("A: Dry Active flower x-axis")),adj=0,cex=1.2)

lines(Rep0dry10,lwd=2,lty=1)
lines(Rep2dry10,col='darkorange1',lwd=2,lty=4)
lines(Rep1dry10,col="dodgerblue",lwd=2,lty=5)
lines(Rep3dry10,col='forestgreen',lwd=2,lty=3)
abline(0,0,lty=3)
segments(-22.,-1,22.,-1,lwd=5,col="black")

#Z
#Generate model lines
Rep0zdry10<-curve(0.40389+((log(x+1))*-0.11189),xlim=c(5,30),ylim=c(0,4))
Rep1zdry10<-curve(0.40389+-0.58718 +((log(x+1))*(-0.11189+0.17614)),xlim=c(5,30),ylim=c(0,4))
Rep2zdry10<-curve(0.40389+-0.58904+((log(x+1))*(-0.11189+0.20061)),xlim=c(5,30),ylim=c(0,4))
Rep3zdry10<-curve(0.40389+-0.59568+((log(x+1))*(-0.11189+0.17110)),xlim=c(5,30),ylim=c(0,4))

#Figure s3C
plot(Change.in.RH.~zoffset,data=Flow10DrymeansREP3Z,xlab="z-axis offset",ylab=expression(paste("",Delta,"RH%")), pch=20,col="forestgreen",ylim=c(-0.2,3),yaxt="n",cex.axis=1.2,cex.lab=1.2)
axis(2,at=c(-2,-1,0,1,2,3,4,5,6,7,8),labels=c("-2","-1","0","1","2","3","4","5","6","7","8"),cex.axis=1.2)

mtext(expression(paste("C: Dry Active flower z-axis")),adj=0,cex=1.2)

points(Change.in.RH.~zoffset,data=Flow10DrymeansREP2Z,pch=20,col="darkorange1")
points(Change.in.RH.~zoffset,data=Flow10DrymeansREP1Z,pch=20,col="dodgerblue")
points(Change.in.RH.~zoffset,data=Flow10DrymeansREP0Z,pch=20,col="black")

lines(Rep0zdry10,lwd=2,lty=1)
lines(Rep1zdry10,col="dodgerblue",lwd=2,lty=5)
lines(Rep2zdry10,col="darkorange1",lwd=2,lty=4)
lines(Rep3zdry10,col="forestgreen",lwd=2,lty=3)
abline(0,0,lty=3)


#Humid Active flowers


#subset data by axis and replicate transect
Flow10WetmeansREP0<-subset(Flow10Wet, subset = sample..=="0")
Flow10WetmeansREP0X<-subset(Flow10WetmeansREP0, subset =xORz==" x")
Flow10WetmeansREP0Z<-subset(Flow10WetmeansREP0, subset =xORz==" z")

Flow10WetmeansREP1<-subset(Flow10Wet, subset = sample..=="1")
Flow10WetmeansREP1X<-subset(Flow10WetmeansREP1, subset =xORz==" x")
Flow10WetmeansREP1Z<-subset(Flow10WetmeansREP1, subset =xORz==" z")

Flow10WetmeansREP2<-subset(Flow10Wet, subset = sample..=="2")
Flow10WetmeansREP2X<-subset(Flow10WetmeansREP2, subset =xORz==" x")
Flow10WetmeansREP2Z<-subset(Flow10WetmeansREP2, subset =xORz==" z")

Flow10WetmeansREP3<-subset(Flow10Wet, subset = sample..=="3")
Flow10WetmeansREP3X<-subset(Flow10WetmeansREP3, subset =xORz==" x")
Flow10WetmeansREP3Z<-subset(Flow10WetmeansREP3, subset =xORz==" z")

#Explore data
#X
#Generate model lines
Rep0Wet10<-curve(3.0736630 +(x*0.0147582)+((x^2)*-0.0033763),xlim=c(-30,30),ylim=c(-8,8))

#Figure s3b
plot(Change.in.RH.~xoffset,data=Flow10WetmeansREP3X,xlab="x-axis offset",ylab=expression(paste("",Delta,"RH%")), pch=20,col="forestgreen",ylim=c(-1,8),yaxt="n",cex.axis=1.2,cex.lab=1.2)
axis(2,at=c(-2,0,2,4,6,8),labels=c("-2","0","2","4","6","8"),cex.axis=1.2)

points(Change.in.RH.~xoffset,data=Flow10WetmeansREP2X,pch=20,col="darkorange1")
points(Change.in.RH.~xoffset,data=Flow10WetmeansREP1X,pch=20,col="dodgerblue")
points(Change.in.RH.~xoffset,data=Flow10WetmeansREP0X,pch=20,col="black")

mtext(expression(paste("B: Humid Active flower x-axis")),adj=0,cex=1.2)

lines(Rep0Wet10,lwd=2,lty=1)
abline(0,0,lty=3)
segments(-22.,-1,22.,-1,lwd=5,col="black")

#Z
#Generate model lines

Rep0zWet10<-curve(2.00937+((log(x+1))*-0.54967),xlim=c(5,30),ylim=c(0,4))

#figure s4b
plot(Change.in.RH.~zoffset,data=Flow10WetmeansREP3Z,xlab="z-axis offset",ylab=expression(paste("",Delta,"RH%")), pch=20,col="forestgreen",ylim=c(-0.2,3),yaxt="n",cex.axis=1.2,cex.lab=1.2)
axis(2,at=c(-2,-1,0,1,2,3,4,5,6,7,8),labels=c("-2","-1","0","1","2","3","4","5","6","7","8"),cex.axis=1.2)

mtext(expression(paste("D: Humid Active flower z-axis")),adj=0,cex=1.2)

points(Change.in.RH.~zoffset,data=Flow10WetmeansREP2Z,pch=20,col="darkorange1")
points(Change.in.RH.~zoffset,data=Flow10WetmeansREP1Z,pch=20,col="dodgerblue")
points(Change.in.RH.~zoffset,data=Flow10WetmeansREP0Z,pch=20,col="black")

lines(Rep0zWet10,lwd=2,lty=1)
abline(0,0,lty=3)

#Seperate figure panels are combined in GNU Image Manipulation Program
#(The GIMP team, GIMP 2.10.8, www.gimp.org) 
```
