## Supplementary file 3 for "Bumblebees can detect floral humidity": Key to Bee Trials Datasheets.docx

Each figure panel (3a, 3b and 4) has two csv file associated with it. First, ‘visit data’ which lists the landing action data for each visit in the respective trials (preferences on active artificial flowers, preferences on passive artificial flowers, and learning). Second, ‘rate data’ that lists humidity response rates over preference tests and success rate over the previous 10 visits in learning trials.

Two ‘.R’ code files are provided: ‘Learning code.R’ and ‘Preference code.R’. These provide sufficient code to repeat analyses for the learning and preference tests respectively. Annotations within the ‘.R’ files are present to guide users through the code. Additionally, code is provided to generate figure 1 and data for figure 2. File ‘Figure 2 gneration.xlsx’ provides a copy of the excel object used to generate figure 2.

In addition to this a list of plotted points for each figure panel is provided in a separate csv file.

Below is a key for each datasheet.

**PassivePreferenceVisitData**

Provides the landing action data for each visit of each of the bees that completed the preference experiment in on the passive artificial flowers.

Column designations are as follows:

bee: bee identifier number within each experiment.

colour: colourings of the bee’s identity marker

date: date of testing

nest: The nest the bee was from. Nests are identified by the lettering indicating the workstation within the Lab their flight area was on. If a nest is replaced during the experiment a number is added to the workstation letter (e.g. D2). Note that nest of the same letter within both active and passive preference tests indicate the same nest, but this is not true of the learning test (see main text).

test group: The test group the bee was in, here the flower type bees were exposed to.

Columns beyond the column labelled ‘visit’ list the foraging actions of each bee. Numbers of columns beyond this point correspond to the flower visit number. Letters correspond to bumblebee foraging actions as follows. ‘H’ landing on a humid flower but no probing. ‘H+’ landing on a humid flower and feeding well is probed. ‘D’ landing on a dry flower but no probing. ‘D+’ landing on a dry flower and feeding well is probed.

**ActivePreferenceVisitData**

Provides the landing action data for each visit of each of the bees that completed the preference experiment in on the active artificial flowers.

Column designations are as follows:

bee: bee identifier number within each experiment.

colour: colourings of the bee’s identity marker

date: date of testing

nest: The nest the bee was from. Nests are identified by the lettering indicating the workstation within the Lab their flight area was on. If a nest is replaced during the experiment a number is added to the workstation letter (e.g. D2). Note that nest of the same letter within both active and passive preference tests indicate the same nest, but this is not true of the learning test (see main text).

test group: The test group the bee was in, here the flower type bees were exposed to.

Columns beyond the column labelled ‘visit’ list the foraging actions of each bee. Numbers of columns beyond this point correspond to the flower visit number. Letters correspond to bumblebee foraging actions as follows. ‘H’ landing on a humid flower but no probing. ‘H+’ landing on a humid flower and feeding well is probed. ‘D’ landing on a dry flower but no probing. ‘D+’ landing on a dry flower and feeding well is probed.

**LearningVisitData**

Provides the landing action data for each visit of each of the bees that completed the learning experiment.

Column designations are as follows:

bee: bee identifier number within each experiment.

colour: colourings of the bee’s identity marker

date: date of testing

nest: The nest the bee was from. Nests are identified by the lettering indicating the workstation within the Lab their flight area was on. If a nest is replaced during the experiment a number is added to the workstation letter (e.g. D2). Note that nest of the same letter are different from those used in the preference tests (see main text).

test group: The test group the bee was in see main text for details. Note that the test group a bee is in dictates which cues correspond with the rewarding and nonrewarding.

Columns beyond the column labelled ‘visit’ list the foraging actions of each bee. Numbers of columns beyond this point correspond to the flower visit number. Letters correspond to bumblebee foraging actions as follows. ‘S’ landing on a rewarding flower but no probing. ‘S+’ landing on a rewarding flower and feeding well is probed. ‘W’ landing on a nonrewarding flower but no probing. ‘W+’ landing on a nonrewarding flower and feeding well is probed.

**PassivePreferenceRateData**

Provides the humidity response rate data (see main text for more information) for each bee that completed the preference test on the Passive artificial flowers.

Column designations are as follows:

bee: bee identifier number within each experiment.

colour: colourings of the bee’s identity marker

date: date of testing

nest: The nest the bee was from. Nests are identified by the lettering indicating the workstation within the Lab their flight area was on. If a nest is replaced during the experiment a number is added to the workstation letter (e.g. D2). Note that nest of the same letter within both active and passive preference tests indicate the same nest, but this is not true of the learning test (see main text).

test group: The test group the bee was in, here the flower type bees were exposed to.

Humid positive actions: The total number (out of 20 visits) of positive responses to humid flower each bee carried out in the preference test (see main test for more detail).

Humidity response rate: The humidity response rate of each bee in the preference test (see main test for more detail).

arcPREF: The arcsine square root transformed humidity response rate, used in analysis.

**ActivePreferenceRateData**

Provides the humidity response rate data (see main text for more information) for each bee that completed the preference test on the Active artificial flowers.

Column designations are as follows:

bee: bee identifier number within each experiment.

colour: colourings of the bee’s identity marker

date: date of testing

nest: The nest the bee was from. Nests are identified by the lettering indicating the workstation within the Lab their flight area was on. If a nest is replaced during the experiment a number is added to the workstation letter (e.g. D2). Note that nest of the same letter within both active and passive preference tests indicate the same nest, but this is not true of the learning test (see main text).

test group: The test group the bee was in, here the flower type bees were exposed to.

Humid positive actions: The total number (out of 20 visits) of positive responses to humid flower each bee carried out in the preference test (see main test for more detail).

Humidity response rate: The humidity response rate of each bee in the preference test (see main test for more detail).

Arcpref: The arcsine square root transformed humidity response rate, used in analysis.

**LearningRateData**

Provides the success rate data calculated at 10 visit intervals in the learning test as described in the main text.

Column designations are as follows:

bee: bee identifier number within each experiment.

colour: colourings of the bee’s identity marker

date: date of testing

nest: The nest the bee was from. Nests are identified by the lettering indicating the workstation within the Lab their flight area was on. If a nest is replaced during the experiment a number is added to the workstation letter (e.g. D2). Note that nest of the same letter are different from those used in the preference tests (see main text).

test group: The test group the bee was in see main text for details. Note that the test group a bee is in dictates which cues correspond with the rewarding and nonrewarding.

visit: the visit number where success rate was calculated: 10, 20, 30, 40, 50, 60, or 70.

Correct actions: the number of correct foraging actions (as defined in the main text) the bee has made in the previous 10 visits.

Success Rate: the success rate achieved by the bee in the previous 10 visits of the learning phase.

ArcCorrect: The arcsine square root transformed success rate, used in analysis.

loglast: The ln transformed Visit, used in analysis.

**Plotted points**

Provides detail on the points plotted in respective figures. For figures 3a and 3b frequencies at different humidity response rates are given. For figure 4 The mean and standard error of the mean of bee success rate is given for each 10 visit division at which the mean was calculated (see main text).
