## Supplementary file 3 for "Bumblebees can detect floral humidity": Learning code.R

```
#Bee learning code from-
#Bumblebees can detect floral humidity

#Authors:
#Michael J. M. Harrap
#Natalie Hempel de Ibarra
#Henry D. Knowles
#Heather M. Whitney
#Sean A. Rands

#Below is the code for the analysis of the bee learning data for the
#aforementioned publication. Instruction and explanation of code is given in
# sections marked with hashtag (#). Outputs from immediately preceding code are given in
#section marked with inverted commas ('').

#For further explanation of the data, statistical models and the model simplification 
#procedure below see the above publication.

#All analysis was carried out in R version 3.6.3 (2020-02-29) -- "Holding the Windsock"

# clear memory and start afresh
rm(list = ls())

#set working directory to file 'Bee trials Data' file
setwd(".../Bee Trials Data") #replace '...' with your directory

#read learning data
Learning<-read.csv(file = "LearningRateData.csv", header = TRUE)

#convert test group to shorthand title (this is done to condense notation of models below)
Learning$test<-Learning$test.group

#Load packages
library(lme4)

#Create seperate datasheets for the different test groups
#This is required to allow analysis of learning within individual
#test groups in the event of an experience~test group interation. 

Con<-subset(Learning, subset = test=="Control")
Dry<-subset(Learning, subset = test=="Dry Reward")
Humid<-subset(Learning, subset = test=="Humid Reward")

#Below we compare effects of
#(ln transformed) experience 'loglost', test group 'test'
#on (asin sqare root transformed) success rate 'ArcCorrect'
#'bee' is included as a random factor

#random intercepts and random slopes model vs just random intercepts
fullmodel<-lmer(ArcCorrect~loglast+test+loglast:test+(loglast|bee), data=Learning, REML = FALSE)
fullmodel2<-lmer(ArcCorrect~loglast+test+loglast:test+(1|bee), data=Learning, REML = FALSE)
anova(fullmodel,fullmodel2)

'Data: Learning
Models:
fullmodel2: ArcCorrect ~ loglast + test + loglast:test + (1 | bee)
fullmodel: ArcCorrect ~ loglast + test + loglast:test + (loglast | bee)
           Df     AIC     BIC logLik deviance  Chisq Chi Df Pr(>Chisq)
fullmodel2  8 -301.29 -271.26 158.64  -317.29                         
fullmodel  10 -298.29 -260.76 159.14  -318.29 1.0029      2     0.6057'

#DeltaAIC
-301.29+298.29
'-3'

#Insufficient difference in AIC.
#standing best model is fullmodel2

#test of interactions interation
fulldropinteration<-lmer(ArcCorrect~loglast+test+(1|bee), data=Learning, REML = FALSE)
anova(fullmodel2,fulldropinteration)

'Data: Learning
Models:
fulldropinteration: ArcCorrect ~ loglast + test + (1 | bee)
fullmodel2: ArcCorrect ~ loglast + test + loglast:test + (1 | bee)
                   Df     AIC     BIC logLik deviance  Chisq Chi Df Pr(>Chisq)    
fulldropinteration  6 -288.16 -265.64 150.08  -300.16                             
fullmodel2          8 -301.29 -271.26 158.64  -317.29 17.125      2  0.0001912 ***'

#DeltaAIC
-301.29+288.16
'-13.13'

#removing interaction raises AIC sufficiently. Best model remains fullmodel2.

#As interaction effects are important, learning effects are analysed at test group level 

#Control group
Conmodel2<-lmer(ArcCorrect~loglast+(1|bee), data=Con, REML = FALSE)
Condroplast<-lmer(ArcCorrect~(1|bee), data=Con, REML = FALSE)
anova(Conmodel2,Condroplast)

'Data: Con
Models:
  Condroplast: ArcCorrect ~ (1 | bee)
Conmodel2: ArcCorrect ~ loglast + (1 | bee)
Df     AIC     BIC logLik deviance Chisq Chi Df Pr(>Chisq)  
Condroplast  3 -121.65 -113.69 63.825  -127.65                          
Conmodel2    4 -124.45 -113.83 66.224  -132.45 4.797      1    0.02851 *'

#DeltaAIC
-121.65+124.45
'2.8'

#insufficient difference in AIC, indicates experience has no effect
#N.B. - fit is improved, perhaps indicates limited experience effect


#Humid Rewards group
Humidmodel2<-lmer(ArcCorrect~loglast+(1|bee), data=Humid, REML = FALSE)
Humiddroplast<-lmer(ArcCorrect~(1|bee), data=Humid, REML = FALSE)
anova(Humidmodel2,Humiddroplast)

'Data: Humid
Models:
Humiddroplast: ArcCorrect ~ (1 | bee)
Humidmodel2: ArcCorrect ~ loglast + (1 | bee)
              Df     AIC     BIC logLik deviance Chisq Chi Df Pr(>Chisq)    
Humiddroplast  3 -79.726 -71.765 42.863  -85.726                            
Humidmodel2    4 -99.966 -89.351 53.983 -107.966 22.24      1  2.406e-06 ***'

#Delta AIC
99.966-79.726 
'20.24'

#removing experience raises AIC sufficiently

#Dry Rewards group
Drymodel2<-lmer(ArcCorrect~loglast+(1|bee), data=Dry, REML = FALSE)
Drydroplast<-lmer(ArcCorrect~(1|bee), data=Dry, REML = FALSE)
anova(Drymodel2,Drydroplast)

'Data: Dry
Models:
Drydroplast: ArcCorrect ~ (1 | bee)
Drymodel2: ArcCorrect ~ loglast + (1 | bee)
Df     AIC     BIC logLik deviance  Chisq Chi Df Pr(>Chisq)    
Drydroplast  3 -36.673 -28.711 21.336  -42.673                             
Drymodel2    4 -74.830 -64.214 41.415  -82.830 40.157      1  2.344e-10 ***'

#Delta AIC
74.830-36.673
'38.157'

#removing experience raises AIC sufficiently


#Figure 2 is generated using Excel, therefore no code is available. However plotted points are given in
#file 'PlottedPoints.csv'.
#File 'Figure 2 generation.xlsx' includes the excel object used for generation of figure 2.
#This excel object undergoes resizing in GNU Image Manipulation Program
#(The GIMP team, GIMP 2.10.8, www.gimp.org) and redundant tick marks for values success rate was not calculated
#at, e.g. 5, 15 etc. on the x axis are removed.

#Below is code that can be used to extract plotted the plotted points.


#Coloum Success.Rate gives means at each visit interval
means<- aggregate(Success.Rate~Visit:test, Learning, function(x) c(mean=mean(x)))

#Coloum Success.Rate gives sd at each visit interval
sd<- aggregate(Success.Rate~Visit:test, Learning, function(x) c(SD=sd(x)))
#Which can be used to calculate SEM as follows
sd$SEM<-sd$Success.Rate/sqrt(15)
```
