## Supplementary file 3 for "Bumblebees can detect floral humidity": Preference code.R

```
#Bee Preference code from-
#Bumblebees can detect floral humidity
  
#Authors:
#Michael J. M. Harrap
#Natalie Hempel de Ibarra
#Henry D. Knowles
#Heather M. Whitney
#Sean A. Rands
  
#Below is the code for the analysis of the bee preference data for the
#aforementioned publication. Instruction and explanation of code is given in
# sections marked with hashtag (#). Outputs from immediately preceding code are given in
#section marked with inverted commas ('').

#For further explanation of the data and analysis see the above publication.

#All analysis was carried out in R version 3.6.3 (2020-02-29) -- "Holding the Windsock"

# clear memory and start afresh
rm(list = ls())

#set working directory to file 'Bee trials Data' file
setwd(".../Bee Trials Data") #replace '...' with your directory


#Passive artificial flowers trial

#read passive preference data
PREF<-read.csv(file = "PassivePreferenceRateData.csv", header = TRUE)


#Below we analyse whether median (asin square root transformed)
#humidity responce rate 'arcPREF', differs from that expected by random foraging

#Expected result of transformed data if random foraging occurs
asin(sqrt(0.5))
'0.7853982'
#further decimals '0.785398163' used below

#Wilcoxon test
wilcox.test(PREF$arcPREF,mu=0.785398163, alternative="two.sided")

'Wilcoxon signed rank test with continuity correction

data:  PREF$arcPREF
V = 109, p-value = 0.005591
alternative hypothesis: true location is not equal to 0.7853982'
#median differs from expected value from random foraging

#Code for fig 1A
hist(PREF$Humidity.response.rate,xlim=c(0.35,0.9),ylim=c(0,6),xlab="Humidity response rate",ylab="Frequency",main=NULL,xaxs="i",yaxs="i",axes=FALSE,breaks=c(0.325,0.375,0.425,0.475,0.525,0.575,0.625,0.675,0.725,0.775))
axis(1,at=c(0.3,0.355,0.4,0.45,0.5,0.55,0.6,0.65,0.7,0.75,0.8,0.85,0.9),labels=c("","",0.4,0.45,0.5,0.55,0.6,0.65,0.7,0.75,0.8,0.85,""),tick=0,las=3)
axis(1,at=c(0.25,0.375,0.425,0.475,0.525,0.575,0.625,0.675,0.725,0.775,0.825,0.875,0.925,0.95,1),labels=c("","","","","","","","","","","","","","",""),tick=-0.01)
axis(2,at=c(0,1,2,3,4,5,6),labels=c(0,1,2,3,4,5,6),las=2)
abline(v=0.5,lty=2,col="black",lwd=2)
box(which="plot", lty="solid")


#_______________________________________

#active artifical flowers trial

#read active preference data
pumpPREF<-read.csv(file = "ActivePreferenceRateData.csv", header = TRUE)

#Below we analyse whether median (asin square root transformed)
#humidity responce rate 'arcPREF', differs from that expected by random foraging

#Wilcoxon test
wilcox.test(pumpPREF$Arcpref,mu=0.785398163, alternative="two.sided")
'Wilcoxon signed rank test with continuity correction

data:  pumpPREF$Arcpref
V = 119, p-value = 0.0008211
alternative hypothesis: true location is not equal to 0.7853982'
#median differs from expected value from random foraging

#Code for fig 1B
hist(pumpPREF$Humidity.response.rate,xlim=c(0.35,0.9),ylim=c(0,6),xlab="Humidity response rate",ylab="Frequency",main=NULL,xaxs="i",yaxs="i",axes=FALSE,breaks=c(0.325,0.375,0.425,0.475,0.525,0.575,0.625,0.675,0.725,0.775,0.825,0.875))
axis(1,at=c(0.3,0.355,0.4,0.45,0.5,0.55,0.6,0.65,0.7,0.75,0.8,0.85,0.9,0.95,1),labels=c("","",0.4,0.45,0.5,0.55,0.6,0.65,0.7,0.75,0.8,0.85,"",0.95,1),tick=0,las=3)
axis(1,at=c(0.25,0.375,0.425,0.475,0.525,0.575,0.625,0.675,0.725,0.775,0.825,0.875,0.925,0.95,1),labels=c("","","","","","","","","","","","","","",""),tick=-0.01)
axis(2,at=c(0,1,2,3,4,5,6),labels=c(0,1,2,3,4,5,6),las=2)
abline(v=0.5,lty=2,col="black",lwd=2)
box(which="plot", lty="solid")


#Seperate panels undergo resizing, croping and are contructed into a single image using
#GNU Image Manipulation Program
#(The GIMP team, GIMP 2.10.8, www.gimp.org)
```
